# Long-term precision editing of neural circuits in mammals using engineered gap junction hemichannels

**DOI:** 10.1101/2021.08.24.457429

**Authors:** Elizabeth Ransey, Kirill Chesnov, Gwenaëlle E. Thomas, Elias Wisdom, Agustin Almoril-Porras, Ryan Bowman, Tatiana Rodriguez, Elise Adamson, Kathryn K. Walder-Christensen, Dalton Hughes, Hannah Schwennesen, Stephen D. Mague, Daniel Colón-Ramos, Rainbo Hultman, Nenad Bursac, Kafui Dzirasa

## Abstract

The coordination of activity between brain cells is a key determinant of neural circuit function; nevertheless, approaches that selectively regulate communication between two distinct cellular components of a mammalian circuit remain sparse. To address this gap, we developed a novel class of gap junctions by selectively engineering two connexin proteins found in *Morone americana* (white perch fish): connexin34.7 (Cx34.7) and connexin35 (Cx35). By iteratively exploiting protein mutagenesis, a novel *in vitro* assay of connexin docking, and computational modeling of connexin hemichannel interactions, we uncovered a pattern of structural motifs that contribute to hemichannel docking compatibility. Targeting these motifs, we designed Cx34.7 and Cx35 hemichannels that dock with each other, but not with themselves, nor with other major connexins expressed in the mammalian central nervous system. We validated these hemichannels *in vivo* using *C. elegans* and mice, demonstrating that they can facilitate communication across neural circuits composed of pairs of distinct cell types and modify behavior accordingly. Thus, we establish a potentially translational approach, ‘<u>L</u>ong-term <u>in</u>tegration of <u>C</u>ircuits using conne<u>x</u>ins’ (LinCx), for context-precise circuit-editing with unprecedented spatiotemporal specificity in mammals.

## Introduction

Many molecular events at synapses regulate the communication between cells, and studies have linked changes in synchrony across brain circuits to cognitive and emotional behaviors. For example, electrical oscillations in hippocampus and prefrontal cortex synchronize during spatial memory in rats (Jones and Wilson, 2005), and between amygdala and hippocampus during fear memory retrieval in mice (Seidenbecher et al., 2003). Moreover, altered long-range synchrony has been observed in preclinical rodent models of schizophrenia (Sigurdsson et al., 2010), depression (Hultman et al., 2018), and autism (Wang et al., 2016), and manipulating synchrony between infralimbic cortex and thalamus has been shown to enhance resilience to acute stress (Carlson et al., 2017). Though these studies highlight the therapeutic potential for modulating synchrony to drive long lasting changes in behavior, it remains a major challenge to manipulate the activity of specific brain circuits with sufficient precision to elicit synchrony. Both spatial (i.e., the interaction of specific cells in space) and temporal constraints of endogenous neural circuits (i.e., the ongoing integrated activity across multiple circuits) make this goal especially challenging. For example, classic pharmacological therapeutics targeting ion receptors or neuromodulatory pathways impact many brain circuits, and emerging brain stimulation-based approaches directly impact the activity of many neuronal populations (i.e., the rate code) and the timing of activity between distinct cells that comprise neural circuits (i.e., the timing code as reflected by synchrony). As such, tools that selectively regulate synchrony within specific circuits in animals have been sparse.

Gap junctions (electrical synapses) enable direct flow of ions and small molecules between two cells and play a prominent role in broadly synchronizing electrical activity in many organs such as the heart and the brain (Alcami and Pereda, 2019; Bruzzone et al., 1996; Laird, 2006). To achieve such synchrony, gap junction consists of two docked segments called hemichannels, embedded in the membranes of two adjoining cells. Each hemichannel, in turn, is an oligomer consisting of six monomeric proteins called connexins (Cx), of which there are 21 isoforms in humans (Sohl and Willecke, 2003; 2004). Most Cxs can form single-isoform hemichannels that dock with themselves to create homotypic gap junctions (Fig. 1A, left).

**Figure 1:**
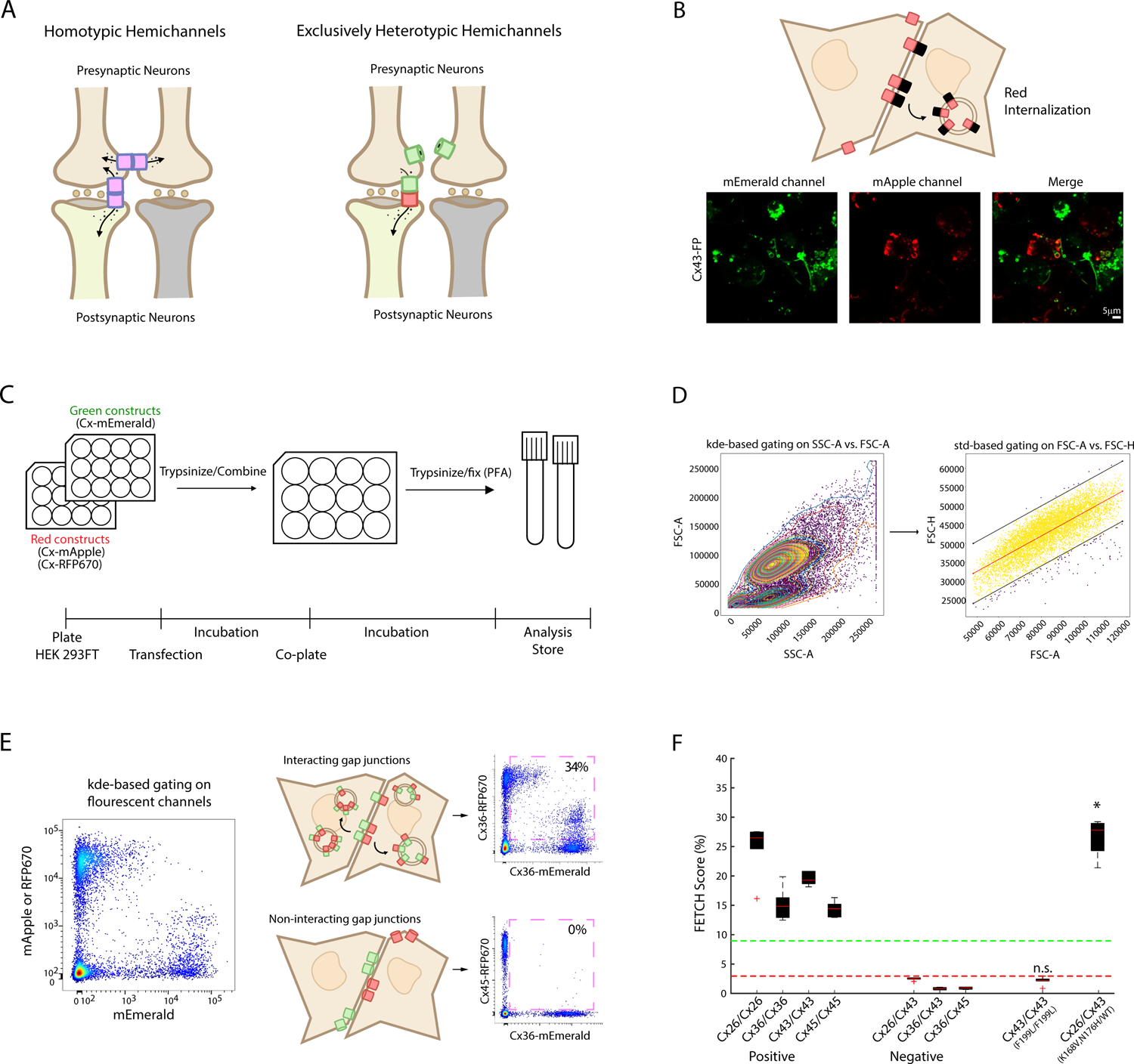
Screen to identify a mutant connexin hemichannel pair that exhibits exclusively heterotypic docking. **A)** Schematic outlining the limitation of introducing ectopic wild type connexin hemichannels (pink rectangles) as a method for ‘selectively’ modulating precise neural circuits composed by brown and light green neurons (left). Note that connexin hemichannels yield off-target electrical synapses between pre-synaptic neurons and, thus, off-target modulation of other circuits. Putative strategy for deploying exclusively heterotypic docking hemichannels (green and red rectangles) to selectively modulate precise neural circuits (right). Note the rectification of this putative gap junction. **B)** Depiction of red fluorescence internalization following gap junction formation (top). Image showing co-plated population of cells expressing counterpart Cx43 tagged to mApple or mEmerald. **C)** FETCH pipeline **D)** Automated gating pipeline that uses sequential kernel density estimate (kde)- and std-based gating approaches **E)** flow cytometry plot obtained from automated pipeline (left). Representative flow cytometry plots for hemichannel pairs with (Cx36/Cx36; top right) and without (Cx36/Cx45; bottom right) docking compatibility. Pink squares highlight portion of transfected cells labeled by two distinct fluorescent proteins. **F)** Portion of dual fluorescent labeled cells for connexin pairs with known docking compatibility profiles (*P<0.05 using t-test compared to distributions of pairs with docking incompatibility).

The exciting potential of employing a gap junction-based tool to modify neural circuit function has previously been established in *C. elegans* (Choi et al., 2020; Hawk et al., 2018; Rabinowitch et al., 2014; Rabinowitch and Schafer, 2015). As with other invertebrates, *C. elegans* do not express connexins. As such, ectopic expression of the vertebrate connexin36 (Cx36) across two connected *C. elegans* neurons results in the formation of an electrical synapse that is inert to endogenous *C. elegans* gap junction proteins (i.e., innexins). Previously work with this approach has successfully been used to modify circuit physiology and multiple behavioral contexts including migration of *C. elegans* in response to various chemical and temperature cues (Choi *et al*., 2020; Hawk *et al*., 2018; Rabinowitch *et al*., 2014; Rabinowitch et al., 2021).

The transformative potential for using gap junctions to repair dysfunctional circuits has been advanced in *C. elegans* as well (Rabinowitch, 2022; Rabinowitch *et al*., 2021). There, circuit editing was utilized to restore normal behavior in animals with circuit impairments. Nevertheless, this work also highlighted a significant challenge to employing gap junctions to edit select circuits in higher-complexity organisms. Specifically, when Cx36 was expressed in two neurons of the same cell class in *C. elegans*, AWC-left and AWC-right, animals showed altered behavior in response to olfactory cues (Rabinowitch *et al*., 2021). Thus, the feature of connexins to form single-isoform homotypic gap junctions dramatically limits its potential to selectively edit brain circuits in vertebrate brains. Here, endogenously expressed Cxs might form gap junctions with the ectopically expressed ones. Moreover, these ectopic connexins might establish undesired connections between individual neurons that may operate in distinct neural circuits but belong to the same cell-type (Fig. 1A, left). As a result, information would flow in an altered manner across many unintended circuits.

Several connexin hemichannels are capable of recognizing hemichannels composed of other connexin protein isoforms to generate heterotypic gap junctions (Fig. 1A, right) (Koval et al., 2014; Laird, 2006). Thus, we reasoned that development and deployment of an exclusively heterotypic hemichannel pairing would provide improved precision in regulating the information flow between distinct cell types in a manner than might be more suitable for use in complex organisms. We reasoned that we could design such a pair by taking advantage of putative mechanisms underlying homotypic and heterotypic hemichannel docking (Bai et al., 2018). We also reasoned that these docking mechanisms, if validated, could be employed to engineer connexin proteins that were docking incompetent to the major connexins endogenous to the mammalian CNS, a feature that would be essential for precise circuit editing in mammalian species.

Goldfish (Carassius auratus) expresses two homologs of mammalian neuronal connexin36 (Cx36) – Cx34.7 and Cx35 – that create a heterotypic gap junction between auditory afferents and Mauthner cells in the CNS (Rash et al., 2013). Interestingly, evidence suggests this Cx34.7/Cx35 heterotypic gap junction rectifies in the Cx34.7 to Cx35 direction (Rash *et al*., 2013). This observation is consistent with the properties of the homologous Cx34.7 and Cx35 proteins found in Morone americana (white perch fish), which also form a heterotypic gap junction that exhibits rectification in the Cx34.7 to Cx35 direction when overexpressed in xenopus oocytes (O’Brien et al., 1998). Moreover, current across this perch gap junction increases above 20mV (O’Brien *et al*., 1998), the threshold for action potential firing in the brains of mammals. Since the rectifying feature of the Cx34.7/Cx35 heterotypic gap junction theoretically enables a preferred directional modulation that approximates at least one important aspect of chemical synapses operation (depolarization of the presynaptic neuron induces depolarization of the postsynaptic neuron, but depolarization of the postsynaptic neuron does not directly induce depolarization of the presynaptic neuron), we chose perch Cx34.7 and Cx35 as our platform to engineer the desired features of a novel synthetic electrical synapse in mammalian cells.

We systematically mutated the residues responsible for Cx34.7 and Cx35 docking to generate two mutant hemichannels that exhibit heterotypic but not homotypic interactions, and do not dock with the major connexin proteins endogenous to the mammalian CNS. We then deployed this synthetic electrical synapse to edit circuit function and behavior in *C. elegans*, recapitulating outcomes observed with an ectopic Cx36/Cx36 gap junction. Finally, we used the engineered Cx34.7/Cx35 synapse in mice across several circuits to demonstrate its functionality and specificity as a neuronal circuit editing tool in mammals.

## Results

### *In vitro* analysis of connexin protein docking

In mammals, connexins are removed from the cell surface via a coordinated endo- and exocytic process resulting in internalization of fully-docked gap junctions within double bi-layer vesicles (Asanuma et al., 1980; Gumpert et al., 2008; Jordan et al., 2001; Mazet et al., 1985; Nickel et al., 2008; Piehl et al., 2007). This feature of connexin regulation has been previously utilized in tissue culture to assess gap junction formation. Specifically, by tagging connexin hemichannels expressed by one cell with a fluorescent protein, internalization into adjacent, non-expressing cells can be visualized (Fig. 1B, top) (Jordan *et al*., 2001; Wang et al., 2015). Since hemichannel docking is a prerequisite for gap junction formation, the trans-internalization of fluorescently-tagged gap junctions can serve as an indicator of hemichannel docking compatibility. In this scenario, fluorescence exchange would be specific for successful hemichannel docking, while diminished fluorescence exchange would be indicative of disrupted hemichannel docking, or alteration of an upstream process critical for expression of connexin hemichannels at the cell surface. Thus, we set out to develop a fast and scalable *in vitro* system to screen the docking compatibility of connexin pairs, utilizing the quantification of fluorescence exchange between cells expressing different fluorescently-tagged hemichannels.

To develop this system, we selected a cell type that rapidly expressed connexin proteins and grew at very high densities that would facilitate surface contacts – a prerequisite for Cx hemichannel docking. Prior work in HEK293FT cells had demonstrated these properties, thus, we expressed individual connexins as either mEmerald-, RFP670-, or mApple fluorescent fusion proteins in HEK293FT cells (Fig. 1B, bottom). Following an initial expression period, separate populations of HEK293FT cells that express intended connexin counterparts were co-plated, incubated, and allowed to grow together overnight. This allows opposed fluorescently-tagged connexins to potentially interact, and the resultant cellular fluorescence exchange was subsequently evaluated via flow cytometry (Fig. 1C-E). Docking is quantified as the portion of transfected cells that are labeled by dual fluorescence in a co-plated sample.

To confirm the functionality of this assay (termed FETCH - <u>f</u>low <u>e</u>nabled <u>t</u>racking of <u>c</u>onnexosomes in <u>H</u>EK cells), we evaluated several connexin proteins that are well established to form homotypic gap junctions that are internalized during normal turnover: Cx26, Cx36, and Cx43 (Forge et al., 2003; Jordan *et al*., 2001; Wang *et al*., 2015). We also evaluated Cx45, which is known to form homotypic gap junctions (Valiunas, 2002) and presumed to be internalized via a similar mechanism. For this evaluation, mEmerald- and RFP670-tagged Cx26, Cx36, Cx43, and Cx45 were transfected into individual populations of cells. When we experimentally paired cells expressing the red- and green tagged Cx isoforms homotypically to determine their fluorescence exchange profiles, we found a substantial portion of dual labeled cells for each Cx (FETCH = 24.8±1.8%, 15.2±1.1%, 19.5±0.4%, and 14.4±0.5% dual-labeled cells for Cx26, Cx36, Cx43, and Cx45, respectively; N=6 replicates per connexin; see Fig. 1F).

We then tested pairs of Cx proteins for which there is evidence of docking-incompatibility, namely Cx26/Cx43, Cx36/Cx43, and Cx36/Cx45 (Koval *et al*., 2014). Here, we determined a substantially smaller proportion of dual-labeled cells (FETCH = 2.5±0.1%, 0.8±0.1%, and 0.9±0.1% for Cx26/Cx43, Cx36/Cx43, and Cx36/Cx45, respectively; N=6 replicates per connexin pair; see Fig. 1F). Critically, the proportion of dual-labeled cells for the population of docking-compatible vs. docking-incompatible pairs was statistically different (T_40_ =14.5; P=1.6×10^-17^ using unpaired t-test), establishing that our *in vitro* screen could be used to broadly assess Cx hemichannel docking compatibility. Finally, we probed whether our FETCH screen could detect the effect of Cx mutations that causally affect gap junction formation. Specifically, we tested a mutation known to disrupt trafficking of Cx43 to the cell membrane, namely F199L (Olbina and Eckhart, 2003), in homotypic configuration. We also evaluated a Cx26 mutant protein, K168V/N176H, known to confer heterotypic docking compatibility with wild-type Cx43 (Karademir et al., 2016). In FETCH experiments, the Cx43-F199L exhibited a level of fluorescence exchange that was statistically indistinguishable from our non-docking heterotypic pairs (FETCH=2.1±0.3%; T_22_ =2.0; P=0.06 compared to pooled distribution of Cx26/Cx43, Cx36/Cx43, and Cx36/Cx45 using unpaired t-test; see Fig. 1F), suggesting a disruption of gap junction formation. On the other hand, the Cx26 mutant/Cx43 wild-type pair exhibited fluorescence exchange that was significantly higher (FETCH=26.6±1.3%; T_22_ =32.7; P=3.9×10^-20^; see Fig. 1F), consistent with docking compatibility. Thus, we established that our FETCH screen was sensitive to connexin mutations that disrupt or enable docking compatibility.

### *In vitro* assessment of Cx34.7 and Cx35 mutant docking

After validating our FETCH assay, we reasoned that this approach could be used to rapidly screen a library of Cx mutations for their ability to disrupt homotypic docking for the two fish Cxs, Cx34.7 and Cx35. Subsequently, a subset of non-homotypic docking Cx34.7 and Cx35 mutant proteins could be screened against each other to identify putative heterotypic docking pairs. Though the precise interactions that guide hemichannel docking are incompletely characterized for the majority of connexins, structure–function and sequence analyses indicate that the second extracellular loop (EL2) plays the greatest role in hemichannel docking specificity (Bai *et al*., 2018; Gong et al., 2013).

To identify potential mutations that disrupt Cx34.7 and Cx35 homotypic hemichannel docking, we rationally introduced seventy and sixty-seven mutations at sixteen positions on both extracellular loops (ELs) of Cx34.7 and Cx35, respectively (see methods for mutant library design; Fig. 2A). We then quantified the strength of homotypic docking interactions of these mutant proteins using FETCH screening (Fig. 2B-C). To identify mutations that completely disrupted docking, we benchmarked mutant homotypic FETCH scores against a heterotypic pairing of human Cx36 and Cx45 proteins that had previously been shown not to yield functional gap junctions (Li et al., 2008). Importantly, for this screening, we were indifferent as to whether these mutations directly disrupted docking interactions or indirectly disrupted docking via interference with an upstream process such as folding or trafficking. Homotypic FETCH screening revealed that most mutants retained their docking property; however, several homotypic non-docking mutant proteins were identified. Homotypic non-docking Cx34.7 mutants included Y78S, Y78T, Y78V, E225K, E225R, L238Y and K222Q, and Cx35 mutants included N56E, Y78V, Y78S, Y78T, E224H, E224K, E224R, and L237Y (Fig. 2B-C)

**Figure 2:**
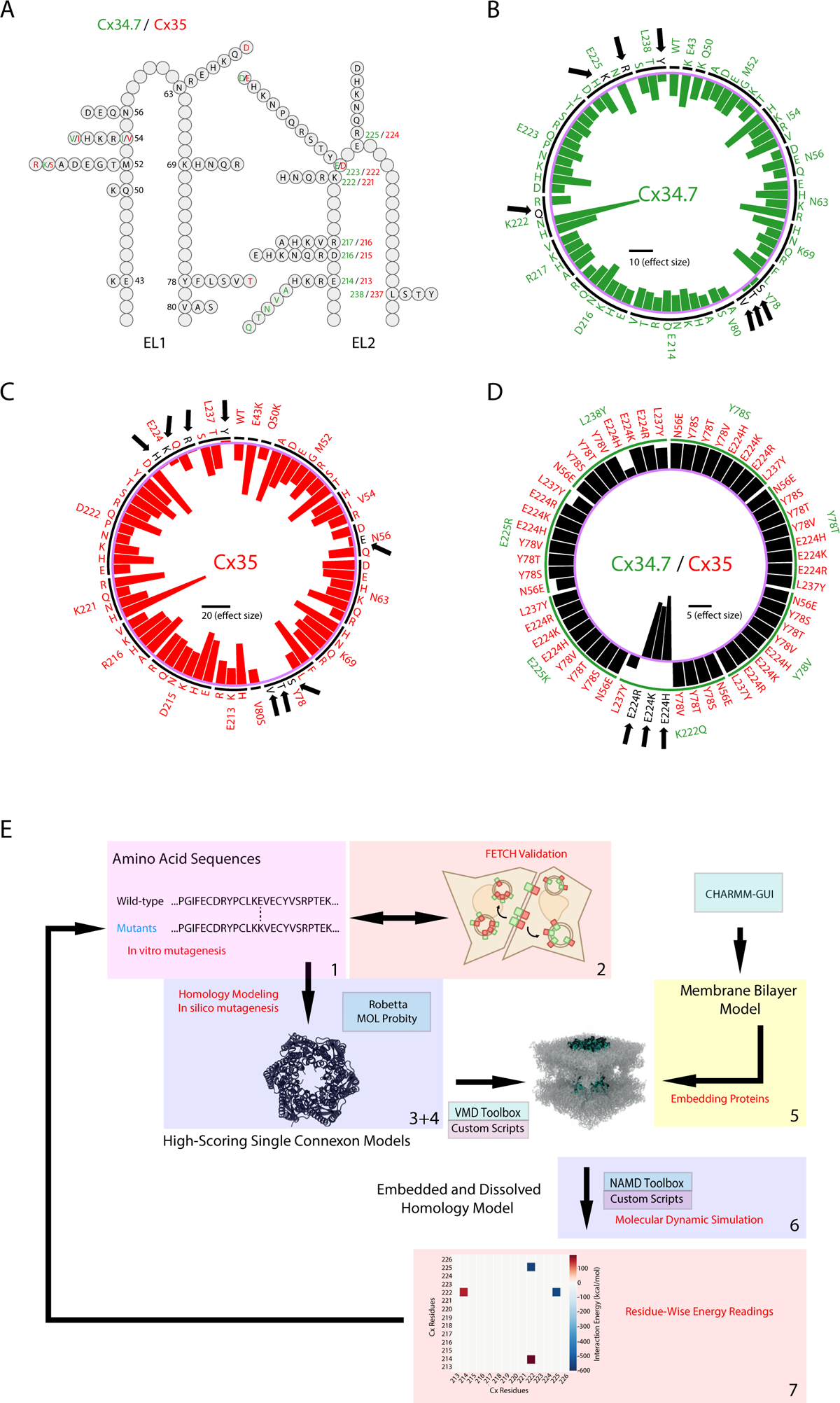
*In vitro* and *in silico* analysis of Cx34.7 and Cx35 mutant docking selectivity. A) Schematic of *Morone Americana* Cx34.7 and Cx35 extracellular loop mutations used to screen for novel, heterotypic exclusive hemichannels. Positions and mutations unique to Cx34.7 and Cx35 are shown in green and red, respectively. Positions and mutations common to both proteins are shown in black. **B-C)** Circular plot showing homotypic FETCH results for B) Cx34.7 and C) Cx35 mutations. Circular bar graphs show the effect size (portion of dual labeled cells) of homotypic mutant combinations relative to the heterotypic pairing of human Cx36 and Cx45 which fails to dock. Black line in the center of the circle corresponds to scale bar for effect size. Targeted residues are listed around the rim of the circle. The substituted amino acids are listed just interior. The intermittent black circle segregates each targeted residue, and the purple circle corresponds to zero effect size. Mutations that disrupted docking are highlighted by black arrows. For example, the black arrow on the left of panel B highlights a mutation from lysine (K) to glutamine (Q) at the 222 resident that disrupted docking of Cx34.7. **D)** Heterotypic FETCH results for Cx34.7 (green) and Cx35 (red) mutant protein combinations. Bar graphs show the effect size of homotypic mutant combinations relative to the wild type Cx34.7 and Cx35 pair. **E)** Integrated approach used to engineer Cx34.7 and Cx35 mutants with docking selectively. Our approach consists of seven integrated components: 1) *in vitro* protein mutagenesis, 2) FETCH screening/validation, 3) homology model generation, 4) *in silico* protein mutagenesis, 5) embedding of proteins in a lipid bilayer and aqueous solution, 6) system minimization, equilibration, and molecular dynamics simulation, 7) and residue-wise energy calculation

Next, to identify mutant protein pairs that exhibit exclusively heterotypic docking, we screened the Cx34.7 and Cx35 homotypic non-docking mutants against each other using FETCH. In this scenario, a substantial portion of dually labeled cells verified that upstream process such as folding/trafficking remained intact for both Cx mutants. We benchmarked these protein pairs against the FETCH scores observed for the wild-type Cx34.7 and Cx35 pair. Strikingly, we discovered three connexin mutant pairs whose FETCH scores were higher than the FETCH scores observed for wild type Cx34.7_WT_/Cx35_WT_ gap junctions. These results provided evidence of mutant pairs (Cx34.7_K222Q_ with either Cx35_E224H_, Cx35_E224K_, or Cx35_E224R_) with heterotypic, but not homotypic, docking (Fig. 2D).

Since our long-term objective was to develop a modulation approach that would be amenable for use in the mammalian nervous system, we also probed whether the four identified mutant isoforms docked with endogenous Cx isoforms expressed in the mammalian brain– specifically Cx36 and connexin43 (Cx43), the major connexins in mammalian neurons and astrocytes, respectively (Condorelli et al., 2000; Rash et al., 2001). Here, we employed FETCH analysis to quantify putative docking, and compared the resultant FETCH scores against established non-docking pair replicates using a one tailed t-test, with a Bonferroni correction for 20 comparisons (92 total non-docking pair replicates; Cx36 and Cx45, FETCH=0.7±0.0%, N=59; homotypic Cx23, FETCH=0.9±0.4%, N=6 replicates; Cx36 and Cx43, FETCH=1.2±0.2%, N=10; and under conditions for which cells were transfected with cytoplasmic fluorophores rather than tagged connexins, FETCH=4.4±0.6%, N=17).

Using this analytical approach, we observed that none of the mutant proteins interacted with human Cx43 (FETCH=1.3±0.1%, 0.4±0.1%, 0.5±0.1%, and 0.5±0.1%; T_96_= 0.29, 1.41, 1.36, and 1.28; P=0.61, 0.92, 0.91 and 0.90 for Cx34.7_K222Q_/Cx43, Cx35_E224H_,/Cx43, Cx35_E224K_/Cx43, and Cx35_E224R_/Cx43, respectively; N=6 replicates for all experimental connexin pairs; FETCH=1.5±0.2%). However, Cx34.7_K222Q_ and the Cx35_E224H_ interacted and formed heterotypic gap junctions with human Cx36 (FETCH=22.8±1.9%; T_91_=-24.4; P=3.4×10^-43^ for Cx34.7_K222Q_/Cx36; FETCH=5.9±1.1%, 0.8±0.1%, and 0.6±0.1%; T_96_=-5.50, 0.89, and 1.24; P=1.6×10^-7^, 0.81, and 0.89 for Cx35_E224H_/Cx36, Cx35_E224K_/Cx36, Cx35_E224R_/Cx36, respectively). Thus, while Cx35_E224K_ and Cx35_E224R_ both showed docking incompatibility with Cx36 and Cx43, and neither showed homotypic docking, we failed to identify an effective Cx34.7 partner that did not dock with Cx36 using our single point mutagenesis and FETCH screening approach.

### Integrating FETCH and *in silico* modeling to design a putatively selective Cx34.7 and Cx35 pair

To address the unintended docking of Cx34.7_K222Q_ with Cx36 we observed via FETCH, we iteratively utilized homology modeling and FETCH analysis to design a new Cx34.7 mutant that does not dock with endogenous Cx43 or Cx36. We concurrently designed its exclusively heterotypic docking Cx35 partner. Briefly, we first developed computational models of the Cx34.7 and Cx35 homotypic and heterotypic gap junctions for wild-type proteins, and several of the mutants that we tested in our initial FETCH screen. We then validated the computational model by comparing the key residues predicted to underly hemichannel docking against the docking characteristics we measured for these mutants using FETCH. Next, to better understand the distinct docking selectivity of the mutant proteins identified in our original FETCH screen (Cx34.7_K222Q_, Cx35_E224H,_ Cx35_E224K,_ and Cx35_E224R_) we modeled their docking interactions with Cx36. Next, we utilized the insights from all our residue-wise interaction models to computationally design Cx34.7 and Cx35 hemichannels that would exhibit exclusively heterotypic docking and would not dock with Cx36 or Cx43. Finally, we generated these proteins and confirmed their docking characteristics *in vitro* using FETCH (Fig. 2E).

First, to model the docking interactions between Cx34.7 and Cx35 gap junctions, we ran molecular dynamics simulations of modeled homotypic and heterotypic pairs of wild type and mutant Cx34.7 and Cx35 proteins (Lee, 2018; Myers et al., 2018). Our molecular dynamic stimulations of the models revealed large negative interaction energies involving residues E214, K222, E223, and E225 in wild type Cx34.7 and residues E213, K221, D222, E224 in wild type Cx35 for both the homotypic and heterotypic docking simulations. These large negative interaction energies were suggestive of salt bridges that stabilize both homotypic and heterotypic docking interactions. These modeling results were consistent with our initial homotypic FETCH screening in which charge swapping mutations (i.e., positive charge to neutral and negative charge to positive at positions Cx34.7-K222 and Cx35-E224, respectively) disrupted docking (see above for FETCH scores). Integrating these results, we identified a common interaction motif for both Cx34.7 and Cx35 consisting of three negative residues (E214/E213, E223/D222, and E225/E224 for Cx34.7 and Cx35, respectively), and a positive residue (K222/K221 for Cx34.7 and Cx35, respectively, see Fig. 3A-C). This interaction motif was consistent with a previously proposed theoretical framework where four residues underlie the docking specificity of most connexin hemichannels (Bai *et al*., 2018).

**Figure 3:**
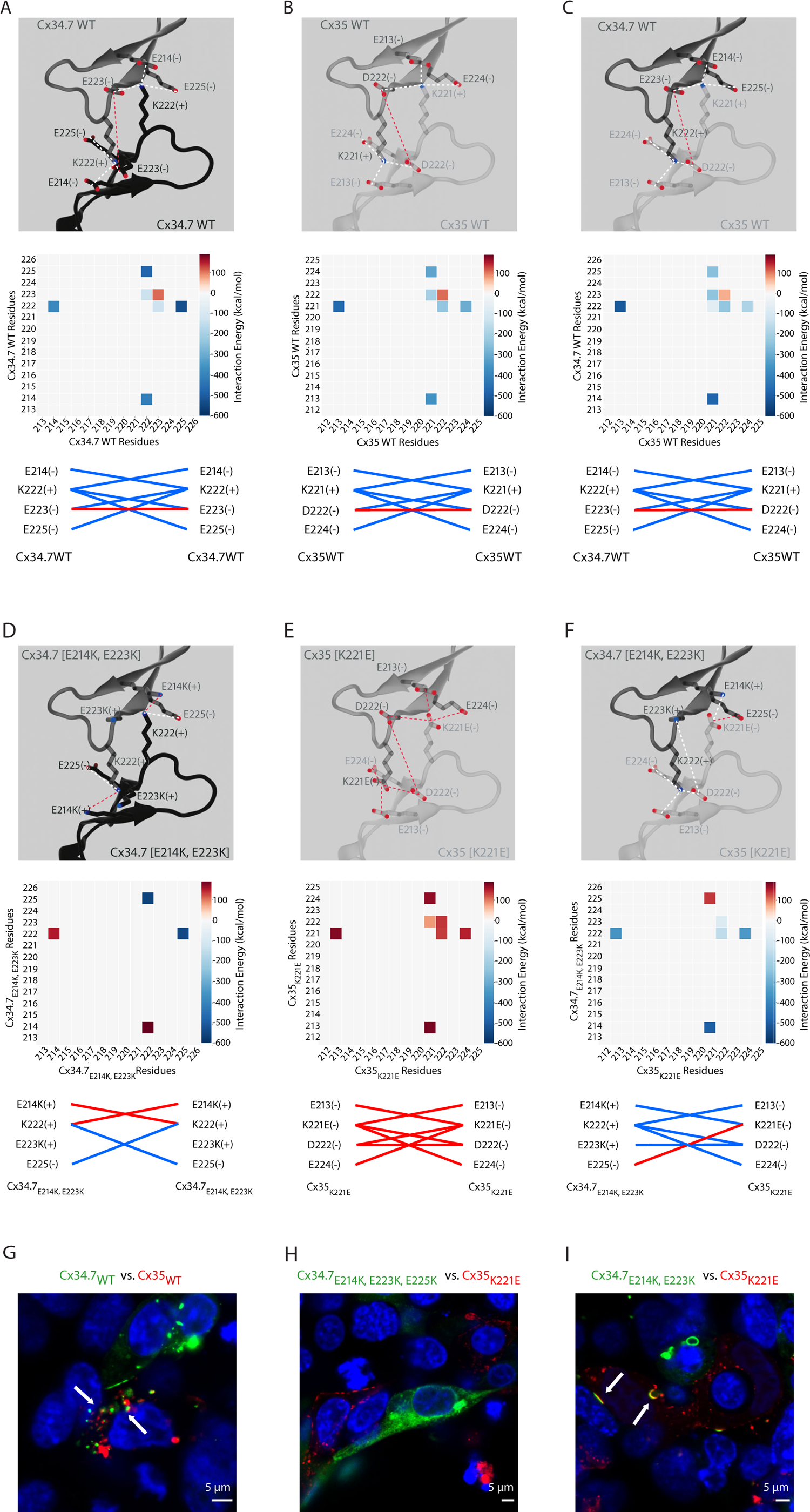
Engineering Cx34.7 and Cx35 mutants to show heterotypic, but not homotypic, hemichannel docking. **A-C)** EL2-to-EL2 interactions predicted between wild type Cx34.7 and Cx35 using homology modeling. Residues predicted to form strong attractive/repulsive interactions are highlighted in blue/red respectively (top). Contact plots for EL2-to-EL2 interactions produced by molecular dynamics simulation (middle), and summary of interactions predicted to stabilize hemichannels pairs (bottom). Plots are shown for A) homotypic Cx34.7, B) homotypic Cx35, and C) heterotypic Cx34.7 and Cx35 interactions. **D-F)** Homology models predicting EL2-to-EL2 residue interactions for Cx34.7 and Cx35 mutant hemichannels. Plots are shown for D) homotypic Cx34.7_E214K, E223K_, E) homotypic Cx35_K221E_, and F) heterotypic Cx34.7_E214K, E223K_ and Cx35_K221E_ interactions (Cx34.7 residues are shown along the y-axis and Cx35 residues are shown along the x axis). **G-I)** Confocal images of heterotypic connexin pairs G) Cx34.7_WT_/Cx35_WT_, H) Cx34.7_E214K, E223K, E225K_/Cx35_K221E_, and I) Cx34.7_E214K, E223K_/Cx35_K221E_ expressed in HEK 293FT cells. All Cx34.7 and Cx35 proteins are expressed as mEmerald and RFP670 fusion proteins, respectively. White arrows highlight dual fluorescent labeled vesicles. Note the cytoplasmic localization of Cx34.7_E214K, E223K, E225K_ in panel H.

Next, we introduced Cx36 into our computational pipeline to generate heterotypic models and residue-wise interaction predictions. Both wild-type Cx34.7 and Cx35 showed strong interactions with Cx36, paralleling the significant FETCH scores we observed (FETCH=11.9±1.2%; T_96_ =-12.93; P=4.7×10^-23^ for Cx34.7/Cx36; FETCH=18.0±2.0%; T_96_=-18.69; P =4.7×10^-34^ for Cx35/Cx36). We then modeled the four non-docking connexin mutants identified in our initial FETCH analysis (Cx34.7_K222Q_, Cx35_E224H_, Cx35_E224K_, Cx35_E224R_) against Cx36. Though the K222Q mutation disrupted the large negative interaction energies we observed in the homotypic wild type Cx34.7 model, the three remaining negative residues within the motif that defines docking compatibility in Cx34.7_K222Q_ continued to show large negative interaction energies with the positive central lysine residue of Cx36, explaining the heterotypic docking between Cx34.7_K222Q_ and Cx36 we observed via FETCH. On the other hand, the three candidate Cx35 mutants we tested against Cx36 using FETCH (Cx35_E224H_, Cx35_E224K_, and Cx35_E224R_) maintained the positive K221 residue that formed strong interactions with the negative residues of Cx36; however, the Cx35_E224K_, and Cx35_E224R_ mutations induced strong repulsion with the positive K238 residue of Cx36, explaining why these two mutants failed to heterotypically dock with Cx36 in our FETCH analyses. Additionally, introducing a smaller positively charged residue at the E224 position, as observed with the Cx35_E224H_ mutant, was sufficient to restore the interaction with Cx36 in the computational model – again mirroring the heterotypic docking profile we observed from our FETCH analyses.

Having modeled the putative interaction principles underlying the docking specificity between Cx34.7, Cx35, and Cx36, and validated our models using FETCH, we set out to design a Cx34.7/Cx35 pair that would exhibit isoform-specific, exclusively heterotypic docking. Our initial strategy was to mutate residues at the four positions of our identified docking motif such that one connexin isoform contained all negatively charged interactors, and the counterpart connexin contributed all positively charged interactors. Our all negatively charged at motif residues mutant, Cx35_K221E_, showed strong repulsions in our homotypic model (Fig. 3E), and did not exhibit homotypic docking upon FETCH analysis (FETCH=1.2±0.4%; T_96_=0.35; P=0.64). Additionally, this mutant protein failed to dock with Cx36 and Cx43 (FETCH=1.5±0.1%, T_91_=0.02, P=0.51 and 1.7±0.2, T_96_=-0.32, P=0.37 for Cx35_K221E_/Cx36 and Cx35_K221E_/Cx43, respectively). Similarly, our all positively charged motif mutant, Cx34.7_E214K/E223K/E225K_, showed strong repulsions in our homotypic computation model and did not exhibit homotypic docking in FETCH analysis (FETCH=0.2±0.0%; T_96_=1.76; P=0.96). However, when Cx35_K221E_ was paired with Cx34.7_E214K, E223K, E225K_ for heterotypic FETCH analysis, the two mutant proteins did not dock (FETCH=1.2±0.3; T_96_=0.37; P=0.64), despite our model predicting strong interaction. Follow-up confocal imaging analysis of HEK 293FT cells expressing the constructs revealed that Cx34.7_E214K, E223K, E225K_ failed to properly localize to the cell membrane (compare Fig. 3G and 3H). We did not evaluate this failed mutant against Cx36 or Cx43; rather, we evaluated an intermediate Cx34.7 mutant protein that maintained positively charged residues at three of the four critical interacting positions, Cx34.7_E214K/E223K_. This mutant showed repulsive interactions in our computational homotypic gap junction model (Fig. 3D), and it showed strong attractive interactions with Cx35_K221E_ (see Fig. 3F). The mutant also localized to the cell membrane where it docked with Cx35_K221E_ as confirmed via FETCH analysis and confocal microscopy (FETCH=35.7±4.1%; T_96_=-28.11; P=2.0×10^-48^; see Fig. 3I). Critically, Cx34.7_E214K/E223K_ did not show homotypic docking in our FETCH analysis (FETCH=1.1±0.2%, T_96_=0.46, P=0.68), nor did it dock with Cx36 or Cx43 (FETCH=1.0±0.2, T_96_=0.58, P=0.72 and FETCH=0.9±0.1%, T_96_=0.73, P=0.77 for Cx34.7_E214K/E223K_/Cx36 and Cx34.7_E214K/E223K_/Cx43, respectively). Curiously, the Cx34.7_E214K/E223K_ and Cx35_K221E_ mutant pair showed a higher heterotypic FETCH score than Cx36 under homotypic docking conditions (FETCH=15.2±1.1%) and the wild type Cx34.7/Cx35 pair (FETCH=12.0±0.9%), as measured using our *in vitro* assay (T_10_=4.9, P=6.4×10^-4^; T_10_=5.7, P=1.9×10^-4^ for comparisons against Cx36/Cx36 and wild type Cx34.7/Cx35, respectively, using t-test with FDR correction; N=6 replicates/group). From here on, we refer to this connexin pair, Cx34.7_E214K/E223K_ and Cx35_K221E_, as Cx34.7_M1_/Cx35_M1_ (designer Cxs version 1.0, from *Morone americana*).

### Synthetic Cx34.7_M1_/Cx35_M1_ electrical synapses regulate behavior in vivo

Using our novel designer Cxs, we set out to determine whether Cx34.7_M1_/Cx35_M1_ could synchronize the activity of distinct neurons that compose a circuit in vivo. We capitalized on *C. elegans* for this analysis because the nematode nervous system is composed of well-characterized circuits of individual cells that regulate behavior. Moreover, several groups have established that selectively expressing Cx36 is sufficient to reconstitute a functional electrical synapse between two connected neurons. Specifically, the presence and function of this resultant Cx36/Cx36 synapse has been confirmed via microscopy (Rabinowitch *et al*., 2014), measurements of synaptic physiology (Rabinowitch *et al*., 2014), calcium imaging (Hawk *et al*., 2018), and behavior (Choi *et al*., 2020; Hawk *et al*., 2018; Rabinowitch, 2022; Rabinowitch *et al*., 2014; Rabinowitch and Schafer, 2015; Rabinowitch *et al*., 2021). Thus, we chose to benchmark our new connexin pair in *C. elegans* against the well characterized Cx36/Cx36 synapse. Specifically, we assessed whether we could induce changes in calcium imaging and behavior with Cx34.7_M1_/Cx35_M1_ in a manner that mirrored Cx36/Cx36.

*C. elegans* do not have an innate temperature preference and can thrive in a broad range of temperatures (Hedgecock and Russell, 1975). However, *C. elegans* trained at a particular temperature in the presence of food will migrate towards that temperature when they are subsequently placed on a temperature gradient without food (Hedgecock and Russell, 1975). This learned behavioral preference is in part mediated by plasticity of the chemical synapse occurring between a thermosensory neuron (called AFD, presynaptic) and an interneuron (called AIY, postsynaptic) (Mori and Ohshima, 1995). Critically, plasticity in the thermosensory neuron AFD can be genetically manipulated to affect transmission to AIY, and to predictably code the otherwise learned behavioral preference (Hawk *et al*., 2018).

As with other invertebrates, *C. elegans* do not express connexins. Thus, ectopic expression of vertebrate connexins result in the formation of electrical synapses that are inert to endogenous gap junction proteins. We have previously shown that ectopic expression of Cx36 could be used to edit the thermotaxis circuit in *C. elegans*, by bypassing the presynaptic plasticity mechanisms between thermosensory neuron AFD and interneuron AIY that contribute to the learned temperature preference (Hawk *et al*., 2018). As such, these circuit-edited animals show a persistent preference for warmer temperatures (Fig. 4A). We therefore used the thermotaxis circuit in *C. elegans* to validate *in vivo* the utility of our engineered gap junction proteins via *de novo* formation of electrical synapses (assessed by calcium imaging) and to recode behavior (assessed by quantitative thermotaxis behavior testing).

**Figure 4.**
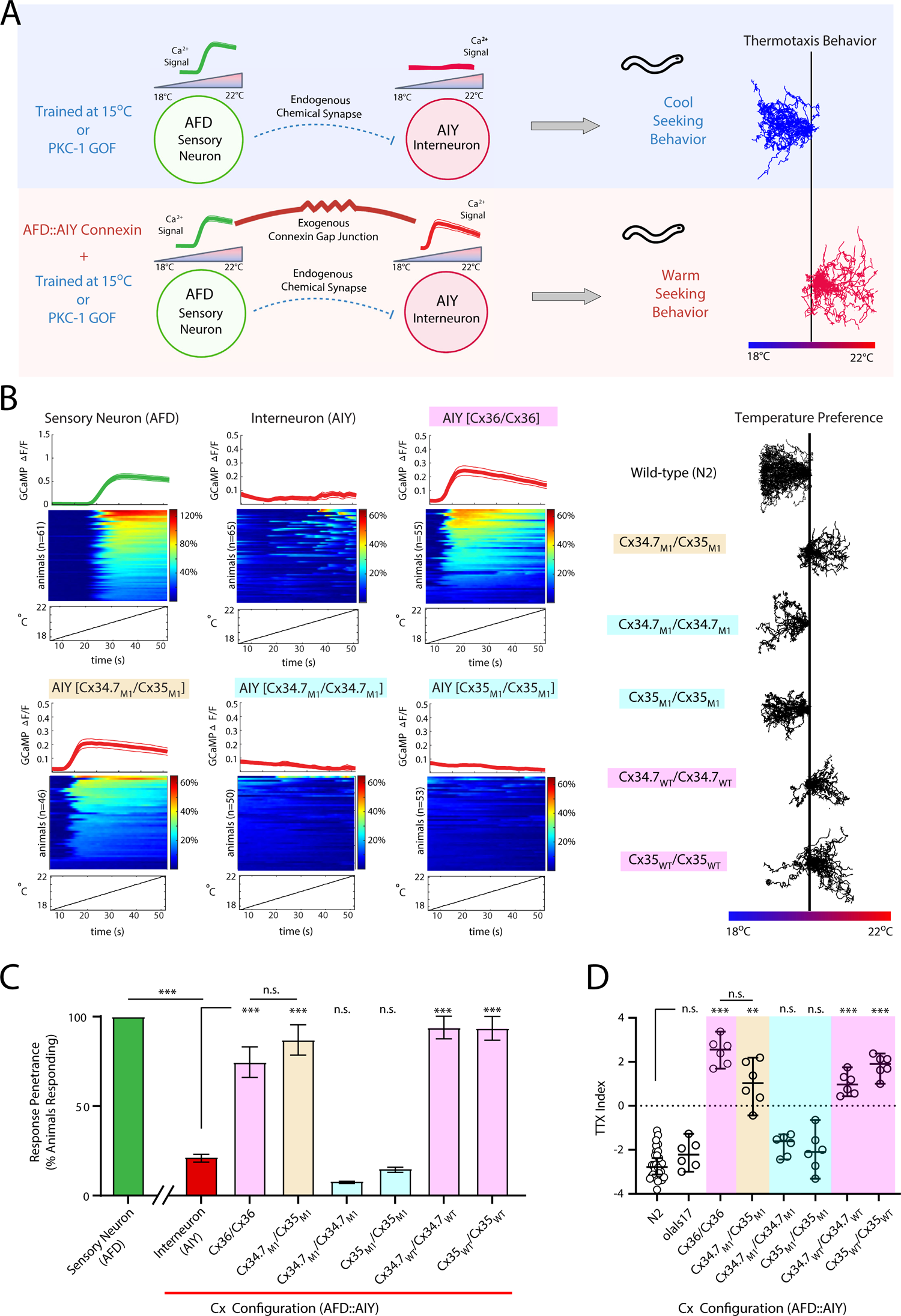
Ectopic connexin hemichannels couple *C. elegans* neurons and recode thermal preference. **A)** Schematic of the AFDàAIY synaptic communication and expressed temperature preference. The AFD thermosensory neuron has a robust calcium response to warming stimuli. *C. elegans* raised in the presence of food at 15°C, or animals with a Protein Kinase C (PKC)*-1* gain-of-function mutation, move towards cooler temperatures when placed on a thermal gradient (top). Ectopic expression of connexin hemichannels between AFD and AIY results in synchronization of the signal to AIY and promotes warm-seeking behavior (bottom). **B)** Calcium traces of neurons expressing ectopic connexin hemichannel pairs (left). Baseline AFD and AIY responses are also shown. Each panel depicts the average trace a group (top, data shown as mean±SEM), heatmaps of individual animals (middle), and the temperature stimulus (bottom). Behavioral traces for each group are shown on the right. Traces are shown for C. *elegans* homotypically expressing wild type connexin hemichannels (pink highlight), heterotypically expressing the mutant pair (tan highlight), and homotypically expressing mutant connexin hemichannels (cyan highlight). **C)** Portion of animals showing neuronal calcium responses based on the traces shown in B; ***p<0.0005 using Fisher’s exact test for penetrance. Error bars denote 95% C.I. **D)** Thermotaxis indices corresponding to experimental groups. Each data point represents the thermotaxis preference index of a separate assay (12-15 animals/assay), with the median for each group plotted denoted by a black horizontal. **p<0.005; ***p<0.0005; Error bars denote 95% C.I.

We first expressed Cx34.7_M1_/Cx35_M1_ cell-specifically in the AFD/AIY pair, respectively, and examined the circuit editing abilities of these proteins (see Supplemental Fig. S1 and Supplemental Table S2). Similar to Cx36/Cx36, expression of Cx34.7_M1_/Cx35_M1_ between the AFD/AIY pair resulted in functional coupling between AFD and AIY, as assessed via calcium imaging (Fig. 4B left and 4C; P<0.0005 using Fisher exact test with an FDR correction). These *C. elegans* constitutively migrated towards warmer temperatures when placed on a thermal gradient, again mirroring the animals expressing exogenous Cx36/Cx36 (F_7,17.91_= 84.99; P<0.0001 using Welch one-way ANOVA followed by Dunnett’s T3 multiple comparisons; p<0.005; Fig. 4B right and 4D). Expression of Cx34.7_M1_ or Cx35_M1_ under homotypic conditions for AFD and AIY neurons (i.e., Cx34.7_M1_/Cx34.7_M1_ or Cx35_M1_/Cx35_M1_) failed to modify circuit function, though homotypic expression for the wild type connexins (i.e., Cx34.7_WT_/Cx34.7_WT_ and Cx35_WT_/Cx35_WT_) synchronized the two cells and modulated behavior (Fig 4B-D; P<0.0005). We also evaluated Cx34.7_M1_ and Cx35_M1_ hemichannels against Cx36 and Cx43 in this live animal model. *C. elegans* expressing Cx34.7_M1_/Cx36, Cx34.7_M1_/Cx43, Cx36/Cx35_M1_ or Cx43/Cx35_M1_ in AFD/AIY all continued to migrate towards cold temperatures (F_7,10.67_= 19.29; P<0.0001 using Welch one-way ANOVA followed by Dunnett’s T3 multiple comparisons; p<0.001; see Supplemental Fig. S3).

Taken together, these findings confirmed that our Cx34.7_M1_/Cx35_M1_ connexin pair modified *C. elegans* behavior and physiology in a manner that was statistically indistinguishable from the well characterized Cx36/Cx36 electrical synapse. Our findings also supported the docking properties we predicted for the mutants using our *in vitro* screen and *in silico* studies, since both Cx34.7_M1_ and Cx35_M1_ failed to alter behavior/physiology when expressed in homotypic configurations or in heterotypic configurations with Cx36 and Cx43.

### Synthetic Cx34.7_M1_/Cx35_M1_ electrical synapse enhances synchrony within a mammalian neural circuit

Having established the *in vitro* docking selectivity and *in vivo* functionality of our Cx34.7_M1_/Cx35_M1_ pair, we set out to determine whether these proteins could modulate neural circuitry in a mammalian species with endogenous connexins. After verifying their expression and trafficking (see Supplemental Fig. S4), we tested whether our proteins could achieve precise circuit editing in mice. Since gap junctions have been shown to play a role in hippocampal function (Allen et al., 2011; Buhl et al., 2003), we compared the impact of expressing either wild-type Cx35 or our Cx35_M1_ mutant in hippocampus. We reasoned that expression of Cx35_WT_, but not Cx35_M1_, would induce hippocampal hyperconnectivity via homotypic docking or heterotypic docking with Cx36 (see Fig. 1B). Thus, we created an adeno-associated virus to target expression of each connexin to neurons (AAV9-CaMKII-Cx35_M1_-mApple and AAV9-CaMKII-Cx35_WT_-mApple), and injected C57BL6/j mice unilaterally in ventral hippocampus (Fig. 5A). We then set out to measure the impact of Cx35_WT_ expression on hippocampal physiology and behavior. Strikingly, expression of Cx35_WT_ was lethal in a dose dependent manner (N=14 mice; Fig. 5B). On the other hand, the Cx35_M1_ mutant was well tolerated (N=16; Fig. 5B). While the mechanism underlying the lethality of Cx35_WT_ in mice remains to be clarified, these results clearly demonstrate the limitation of Cx35_WT_ for use in mammalian circuit editing.

**Figure 5.**
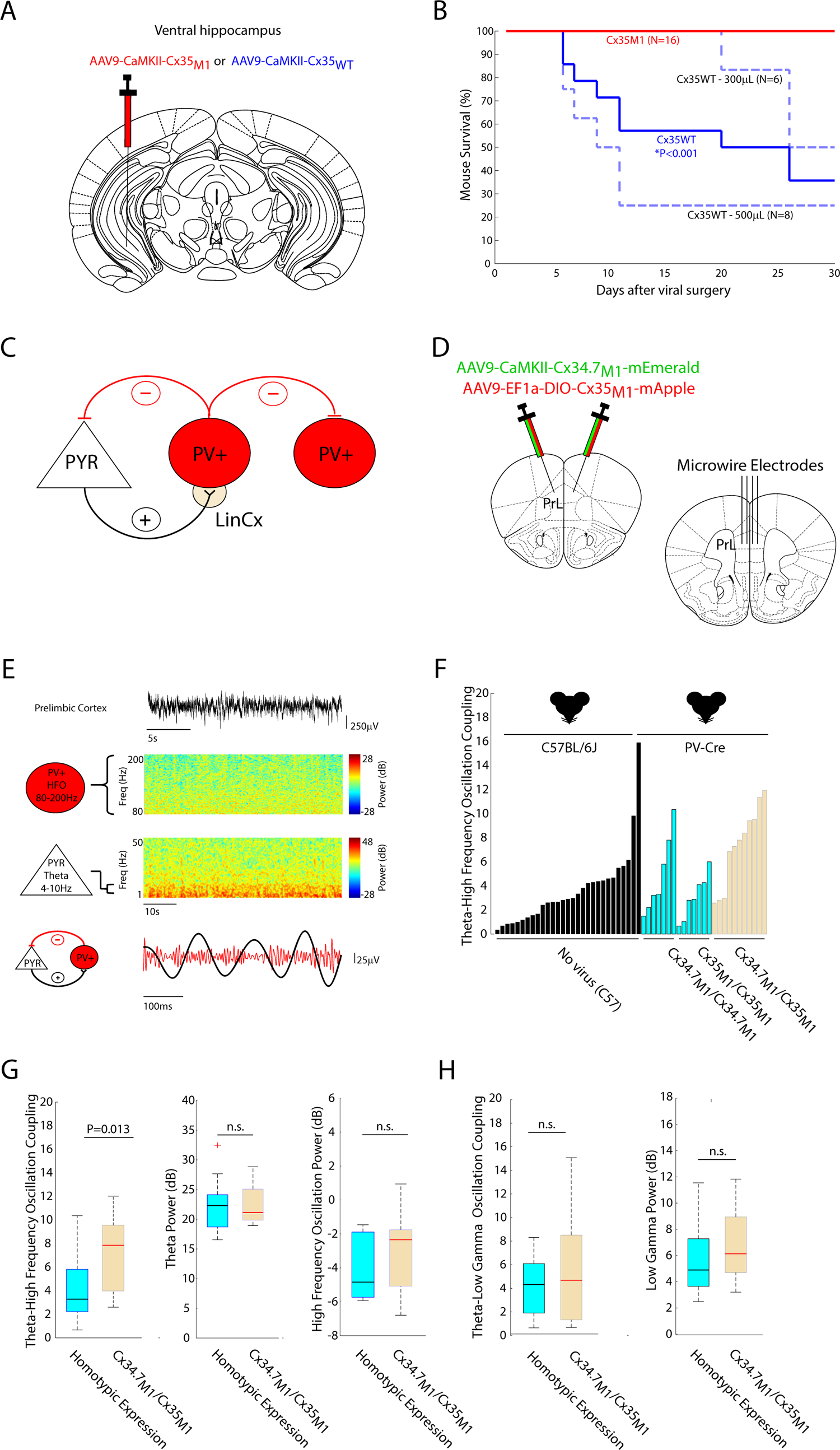
LinCx edits microcircuit dynamics at the millisecond timescale in mice. **A)** Mice were injected with AAV-CaMKII-CX35_WT_ or with AAV-CaMKII-CX35_M1_ in ventral hippocampus. **B)** Survival curves for Cx35_WT_ vs. CX35_M1_ injected mice. **C)** Prefrontal cortex microcircuit comprising an excitatory pyramidal neuron (PYR) and a parvalbumin expressing fast-spiking interneuron (PV+). The tan circle highlights the target for synaptic editing (top). **D)** PV-Cre mice were bilaterally co-injected with AAV-CaMKII-Cx34.7_M1_ and AAV-DIO-Cx35_M1_ (bottom). Control mice were injected with a Cx34.7_M1_ or Cx35_M1_ pair of viruses which expressed the same hemichannel in both cell types. Mice were subsequently implanted with microwires in prefrontal cortex. **E)** Representative local field potential recorded from prefrontal cortex (top). Power spectrograms show theta (4-10Hz) and high frequency activity (80-200Hz), which corresponds to PYR and PV+ neuron firing, respectively (middle). Microcircuit function is represented by the coupling between the phase of theta oscillations (black) and the amplitude of high frequency oscillations (red, bottom). **F)** Distribution of theta–high frequency coupling scores observed across un-injected C57BL/6j mice (N=29, black bars), PV-Cre mice co-injected with a Cx34.7_M1_ (N=7) or Cx35_M1_ (N=7) pair of viruses (blue bars), or mice co-injected with a docking Cx34.7_M1_/Cx35_M1_ pair (N=11, tan bars). **G)** Mice co-injected with the Cx34.7_M1_/Cx35_M1_ pair showed significantly higher theta-high frequency oscillation (left), but not theta power or high frequency oscillation power (middle, right), compared to mice expressing Cx34.7_M1_ and Cx35_M1_ under homotypic conditions expressing. **H)** Circuit editing had no impact on theta-low gamma oscillatory coupling (left) or low gamma power (right).

After establishing the potential of the mutant proteins for mammalian circuit editing, we chose to edit a neurocircuit composed of two distinct cell types. We used mice to achieve cell type specificity, because they are highly amenable to cell-type specific access via selective promoters and Cre-recombinase targeting. Excitatory pyramidal neurons (PYR) and parvalbumin expressing fast-spiking interneurons (PV+) can form microcircuits whereby PYR neurons excite PV+ neurons, which in turn inhibit PYR neurons (Fig. 5C). This PYR↔PV+ neural circuit has been well characterized in the hippocampus, where PYR neurons show activity coupled to the phase of theta frequency (4-10Hz) oscillations during spatial exploration (Siapas et al., 2005) and PV+ neuron activity correlates with gamma frequency oscillations (30-80Hz)(Fuchs et al., 2007). Critically, the activity of this PYR↔PV+ microcircuit is reflected in the millisecond resolved synchrony between the phase of theta oscillations and the amplitude of gamma oscillations in rodents (Wulff et al., 2009).

PYR↔PV+ microcircuits are also observed in the prefrontal cortex with slightly different neurophysiological properties (Sohal et al., 2009). Like the cellular dynamics observed in the hippocampus, prefrontal cortex PYR neurons phase-couple to locally recorded theta oscillations (Dzirasa et al., 2010). On the other hand, in prefrontal cortex, PV+ neurons best couple to the phase and amplitude of local high frequency oscillations (80-200Hz)(Yao et al., 2020). Thus, to determine the effect of our synthetic synapses, we quantified the coupling between the phase of prefrontal cortex theta oscillations and the amplitude of prefrontal cortex high frequency oscillations as a proxy for prefrontal cortex PRY ↔ PV+ microcircuit activity. Specifically, we expressed our synthetic Cx34.7_M1_/Cx35_M1_ synapse at the PRY→PV+ interface, hypothesizing that this manipulation would enhance the millisecond-timed coupling between theta and high frequency oscillations in the prefrontal cortex.

Specifically, we developed an AAV (AAV9-CaMKII-Cx34.7_M1_-mEmerald) to express Cx34.7_M1_ in PYR neurons, and another virus (AAV9-Ef1α-DIO-Cx35_M1_-mApple) to target Cx35_M1_ to cells expressing Cre-recombinase. We then co-injected PV-Cre mice with both viruses bilaterally in the prelimbic cortex (PrL, a subdivision of prefrontal cortex in mice) such that Cx34.7_M1_ is expressed in PYR neurons (though non-selectively) and Cx35_M1_ is selectively expressed in PV+ neurons (i.e., PYRàPV+ modulation given the putative Cx34.7àCx35 rectification of our hemichannel pair; N=11; Fig. 5C-D). A control distribution of non-edited mice consisted of three groups: C57BL/6J mice that had not been injected with virus (N=29), PV-Cre mice co-injected with AAV9-CaMKII-Cx34.7_M1_-mEmerald and AAV9-Ef1α-DIO-Cx34.7_M1_-mEmerald (which expressed Cx34.7_M1_ in the homotypic non-docking configuration; N=7), and PV-Cre mice co-injected with AAV9-CaMKII-Cx35_M1_-mApple and AAV9-Ef1α-DIO-Cx35_M1_-mApple (to express Cx35_M1_ in the homotypic non-docking configuration; N=7). All mice were subsequently implanted with microwire electrodes bilaterally in prelimbic cortex (see Supplemental Fig. S5A). Neural oscillatory activity was recorded while mice explored an open field.

To determine the coupling between theta (4-10Hz) and high frequency oscillations (80-200Hz), we isolated local field potential activity in these two frequency bands (Fig. 5E). We then determined their phase–amplitude relationships using the established modulation index (z-score), which quantifies the statistical likelihood that measured relationships between two oscillations would be observed by chance (Canolty et al., 2006). Using this approach, we found significant theta–high frequency oscillation coupling from the majority implanted mice (80%, 43/54; Fig. 5F). Moreover, theta–high frequency oscillation coupling was significantly higher in the mice expressing our synthetic synapse, compared to the pooled group of control mice expressing the channels under homotypic configurations (U=141; P<0.013 using one-tailed rank-sum test). Thus, we found that expression of the synthetic synapse was sufficient to enhance millisecond-timed synchrony within a circuit defined by two precise cell types in mammals. Interestingly, our post-hoc analysis also found no differences in theta–high frequency oscillation coupling between mice expressing Cx34.7_M1_/Cx34.7_M1_ or Cx35_M1_/Cx35_M1_ in the homotypic non-docking configuration compared to uninjected C57BL/6J control mice (U=509 and 542; P=0.28 and 0.84 using two tailed rank-sum test, for Cx34.7_M1_/Cx34.7_M1_ or Cx35_M1_/Cx35_M1_, respectively). These findings from our homotypic controls support the non-homotypic docking selectivity of the two mutant proteins.

Importantly, no differences in theta or high frequency oscillation power were observed between mice expressing the synthetic synapse and the control mice expressing mutant connexins in homotypic configurations across the PYRàPV+ circuit (U= 172 and 167; P=0.60 and 0.43 using two tailed rank-sum test, for theta and high frequency oscillation power, respectively; Fig. 5G). Similarly, no differences in theta oscillations and low gamma oscillations cross frequency phase coupling were observed across these groups (U= 176, P=0.76 using two tailed rank-sum test; Fig. 5H). Thus, when expressed in the prelimbic cortex PYRàPV+ circuit, the synthetic synapse selectively regulated synchrony between theta and high frequency oscillation activity.

Since prior evidence suggested that the heterotypic Cx34.7/Cx35 gap junction rectifies in the Cx34.7 direction (O’Brien *et al*., 1998; Rash *et al*., 2013), we wondered whether reversing the orientation of the synthetic synapse between PRY and PV+ would also impact synchrony in the circuit. Thus, we developed an adeno-associated virus (AAV9-EF1α-DIO-Cx34.7_M1_-mEmerald) to selectively target Cx34.7_M1_ to cells expressing Cre-recombinase, and another virus (AAV9-CAMKII-Cx35_M1_-mApple) to express Cx35_M1_ in PYR neurons. Again, we infected PV-Cre mice with these viruses and implanted recording electrodes in PrL (N=5 mice). When we compared theta-high frequency oscillation coupling in these mice to new control mice infected with fluorescent proteins viruses (N=13; AAV9-EF1α-DIO-mCherry and AAV9-CaMKII-GFP; N=15 mice), we found no difference between groups (U= 125, P=0.92 using two tailed rank-sum test; see Supplemental Fig. S6). This suggests that the impact of our Cx34.7_M1_ /Cx35_M1_ gap junction on circuit function is at least in part determined by the orientation of the Cx35_M1_ and Cx34.7_M1_ hemichannels, providing evidence to support rectification of the channel and underscoring the possible use of the engineered proteins to achieve directional excitability between target neurons.

### Synthetic Cx34.7_M1_/Cx35_M1_ electrical synapse potentiates a multi-regional circuit in mammals

Having established that our synthetic electrical synapse could enhance synchrony within a local microcircuit, we next tested whether it could potentiate a long-range circuit. The Infralimbic cortex (IL, another anatomical subdivision of mouse medial prefrontal cortex) sends a monosynaptic projection to medial dorsal thalamus (MD) in mice. We selected this circuit to characterize the impact of LinCx on long-range circuity given our prior experience in quantifying the role of cortico-thalamic circuitry in physiology and behavior (Carlson *et al*., 2017; Kumar et al., 2013). Specifically, in our prior work, we found that a direct optogenetic pulse to medial prefrontal cortex induced a positive potential in medial dorsal thalamus within 25ms (Kumar *et al*., 2013), likely due to cortical activation of local inhibitory networks. Thus, we hypothesized that expression of our synthetic synapse between IL terminals and cell bodies in MD would strengthen the thalamic potential that was induced by cortical stimulation.

To target this circuit, we co-injected BALB/cJ mice with AAV9-CaMKII-Cx34.7_M1_-mEmerald and AAV9-CamKII-Chr2-EYFP in left IL. Three weeks later, we injected these mice with AAV9-CaMKII-Cx35_M1_-mApple in left MD (i.e., ILàMD modulation, given Cx34.7àCx35 rectification of the hemichannel pair) and implanted microwire recording electrodes in IL and MD (Fig. 6A). After another 5 days of surgical recovery, we recorded LFP activity from both regions in response to 10ms pulses of optogenetic stimulation to IL during free behavior. This timeline ensured expression of Chr2 and Cx34.7_M1_ in IL, but minimal trafficking of Cx34.7_M1_ to the IL axonal terminals in MD (see Supplemental Fig. S4), and minimal local expression of Cx35_M1_ in MD. We acquired additional recording and stimulation data nine days later (5 weeks after the initial IL injection/ 2 weeks after the MD injection), enabling strong trafficking of Cx34.7_M1_ to IL and substantial local Cx35_M1_ expression. A control group was infected with AAV9-CaMKII-GFP in MD instead of Cx35_M1_.

**Figure 6.**
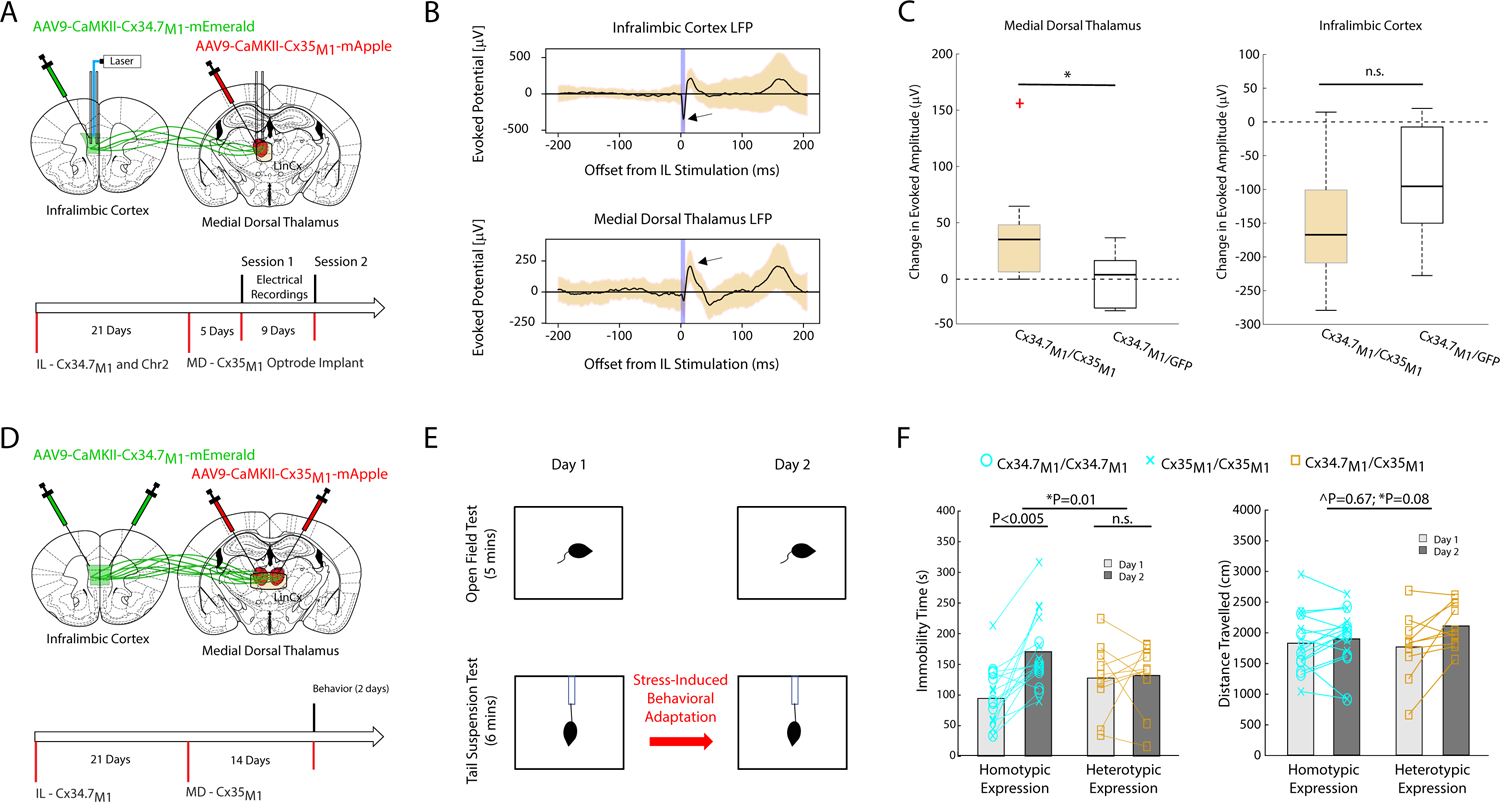
LinCx edits microcircuit dynamics at the millisecond timescale in mice. **A)** Schematic of viral and opto-electrode targeting approach (top), and experimental timeline for optogenetic interrogation of the ILàMD circuit (bottom). **B)** Representative light evoked response recorded from IL (top) or MD microwire (bottom). Light stimulation was delivered at 1mW, 10ms pulse width. Data shown as mean ± std across light pulses. **C)** Change in amplitude of evoked potential across sessions in MD (left) and IL (right). *P<0.05 using one tailed t-test. **D)** Schematic of viral injection strategy and experimental timeline for quantifying impact of ILàMD LinCx editing on behavior. **E)** Schematic of behavioral testing to quantify stress induced behavioral adaptation. **F)** Immobility time and distance travelled during repeat tail suspension (left) and open field testing (right). Mice co-injected with Cx34.7_M1_ (N=8) or Cx35_M1_ (N=8) in homotypic non-docking configurations showed stress-induced behavioral adaptation during repeat TST testing (blue), while mice co-injected with the functional Cx34.7_M1_/Cx35_M1_ pair did not (N=10; tan). No significant behavioral differences were observed between viral groups in the open field; ^ denotes Group effect; * denotes Group × Day interaction effect using mixed effects model ANOVA.

Consistent with our prior study, we observed a positive evoked potential in MD within 25ms of IL stimulation with 1mW of blue light during the first session (Fig. 6B, see also Supplemental Fig. S7). When we repeated our stimulation experiment nine days later, almost all the mice expressing Cx35_M1_ showed an increase in the amplitude of their evoked MD activity (N=8/9 mice, 41±16mV). This increase was significantly higher than what we observed from the control group across sessions (N=6, 1±10mV; t_13_=1.9, P= 0.043 using one tailed t-test; see Fig. 6C, left). There was no statistical difference between the change in the evoked IL response across groups (−154±30mV and −93±38mV for the Cx35M1 and GFP groups, respectively; t_13_=-1.3, P= 0.11 using one tailed t-test; see Fig. 6C, right). Taken together, these findings showed that expression of our synthetic synapse potentiated the ILàMD circuit.

### Synthetic Cx34.7_M1_/Cx35_M1_ electrical synapse modifies behavior in mammals

Finally, we set out to determine whether our synthetic electrical synapse could be used to modify behavior. Here, we again targeted the ILàMD circuit. Our prior work had implicated this circuit in mediating stress resilience. The tail suspension test is a classic assay that measures the behavioral response of mice to an inescapable negative experience in which they are suspended upside down by their tail (Steru et al., 1985). Prior exposure to stress diminishes behavioral responses during the assay (Iniguez et al., 2016), and the assay itself induces a robust stress response (Ide et al., 2010). We previously exploited these two features of the test to demonstrate that repeated exposure to the tail suspension test increases immobility during subsequent testing. Critically, no increases in immobility are observed when exploratory behavior is assayed immediately prior to the tail suspension sessions; demonstrating that the behavioral adaptation is specific to the stressful context (Carlson *et al*., 2017). The stress-induced behavioral adaptation observed following exposure to the tail suspension test is accompanied by compensatory changes in IL and MD, and optogenetic stimulation of the ILàMD circuit, in a manner that recapitulates the normal synchrony in the circuit, and reduces the behavioral adaptation induced by acute stress (Carlson *et al*., 2017).

Here, we employed this behavioral assay to test whether expression of the synthetic Cx34.7_M1_/Cx35_M1_ electrical synapse across the ILàMD circuit could diminish stress adaptation. Since this manipulation had potentiated the ILà MD compensatory circuit, our hypothesis was that it would prevent the immobility adaptation seen between the two sessions of the tail suspension test. To test this hypothesis, we injected BALB/cJ mice with AAV9-CaMKII-Cx34.7_M1_-mEmerald bilaterally in IL. Three weeks later, we injected them with AAV9-CaMKII-Cx35_M1_-mApple bilaterally in MD (Fig. 6D; N=10 mice; see supplemental Fig. S5B). A negative control group of mice was injected with either AAV9-CaMKII-Cx34.7_M1_-mEmerald (N=8) or AAV9-CaMKII-Cx35_M1_-mApple (N=8) in both regions to express synthetic hemichannels across the ILàMD circuit in homotypic non-docking configurations. Two weeks later, all mice were subjected to two days of testing in an open field and tail suspension (Fig. 6E).

As hypothesized, mice expressing the synthetic synapse across the IL-MD circuit did not show behavioral adaptation in response to repeat tail suspension testing (F_1,24_=7.85, P=0.01 for Group × Day interaction effect using mixed effects model ANOVA; t_9_=0.19; P=0.85 for post-hoc testing using two-tailed paired t-test for Cx34.7_M1_/Cx35_M1_ mice across days; Fig. 6F, left). On the other hand, increases in immobility were observed in the negative control group (t_18_=4.9; P=1.7×10^-4^ for post-hoc testing using two-tailed paired t-test for pooled group of Cx34.7_M1_/Cx34.7_M1_ and Cx35_M1_/Cx35_M1_ mice across days). As anticipated, post-hoc analysis revealed increases in immobility in both the Cx34.7_M1_/Cx34.7_M1_ and Cx35_M1_/Cx35_M1_ control groups expressing synthetic hemichannels in non-docking configurations, independently (t_7_=3.3; P=0.01, and t_7_=5.5; P=9.5×10^-4^, respectively using two-tailed paired t-test). Moreover, these control mice showed increases in TST immobility that were statistically indistinguishable from that observed in uninfected BALB/cJ mice (Supplementary Fig. S8). Thus, expression of our functional synthetic synapse across the ILàMD circuit prevented behavioral adaptation to the tail suspension test. No differences in open field exploration were observed across groups between the mice that expressed Cx34.7_M1_/Cx35_M1_ and control mice expressing the hemichannels in non-docking configurations (F_1,24_=0.19, P=0.67 for Group effect; F_1,24_=3.4, P=0.08 for Group × Day interaction effect using mixed effects model ANOVA), demonstrating that editing ILàMD only changed behavior in the stressful assay (Fig. 6F, right). Both groups of mice displayed higher exploration in the open field during the second testing session (F_1,24_=7.69, P=0.01 for Day effect using mixed effects model ANOVA), demonstrating that the increased immobility displayed by control animals on day 2 of the tail suspension assay was specific to the stress context and not a change in general locomotor activity.

## Discussion

We developed a novel approach we term ‘<u>L</u>ong-term <u>in</u>tegration of <u>C</u>ircuits using Conne<u>x</u>ins’ (LinCx) that can be used to edit brain circuits in mammals. This approach employs an engineered connexin hemichannel pair capable of heterotypic, but not homotypic, docking. When expressed in two adjacent cells, these hemichannels compose a heterotypic electrical synapse, facilitating the transfer of activity between them. We chose to engineer LinCx using two connexin proteins found in *Morone americana* (white perch fish): connexin34.7 (Cx34.7) and connexin35 (Cx35). Our approach is analogous to the strategies for circuit editing previously employed in *C. elegans* using Cx36 (Choi *et al*., 2020; Hawk *et al*., 2018; Rabinowitch *et al*., 2014; Rabinowitch and Schafer, 2015). Indeed, we found that both Cx34.7_WT_ and Cx35_WT_ could be used to edit AFD/AIY under homotypic conditions, yielding physiological and behavioral changes that were equally as robust as Cx36. On the other hand, our engineered proteins, Cx34.7_M1_ and Cx35_M1_ only dock heterotypically. This feature ensures that even when a hemichannel is expressed across a cell type, those cells will not form electrical synapses between themselves (Fig. 1A). This was consistent with our observations in *C. elegans* that found no behavioral or physiological changes when our mutant proteins were deployed to AFD/AIY homotypically. Thus, our LinCx system can be deployed in higher order animals, which have many more cells of each given cell type, to target distinct neural circuits. We demonstrated this novel feature in mice by editing a cortical microcircuit composed of two genetically defined cell types.

To facilitate such strong docking specificity, we employed a three-part strategy to engineer a Cx34.7 and Cx35 pair that solely exhibit heterotypic docking. First, we created a novel approach to rapidly assess the docking of connexin hemichannels. This approach was built on well-established methods utilizing fluorescence exchange between cells expressing fluorescently-labeled connexin hemichannels as an indicator of gap junction formation. The FETCH score as assessed using flow-cytometry then indicates the number of cells within a population that clearly exhibit fluorescence exchange. We validated this approach using pairs of connexin proteins with established docking compatibilities. We then created a library of Cx34.7 and Cx35 mutants based on sites previously shown to confer docking specificity, as well as those implicated in docking based on homology modeling from the structures of Cx26 (Bennett et al., 2016; Maeda et al., 2009). Next, we subjected these mutants to our novel rapid screening approach to identify the mutations that disrupted homotypic docking for each connexin protein. We benchmarked the mutants against a distribution of FETCH scores that was highly specific for, but not necessarily sensitive to, docking incompatibility. While this approach increased our confidence that the mutations we selected for further screening impaired homotypic docking, it remained plausible that several excluded mutations disrupted homotypic docking as well. Next, we used FECTH to screen heterotypic mutant pairs of Cx34.7 and Cx35 against a distribution of FETCH scores that was highly specific for heterotypic docking comparability, namely wild-type Cx34.7 and Cx35 pairs. This strategy increased our confidence that the mutant pairs selected for further analysis exhibited heterotypic docking, though it did not ensure that excluded mutant pairs exhibited heterotypic docking incompatibility. Finally, we characterized the docking specificity of our candidate proteins against human Cx36 and Cx43. Here, our goal was to identify candidates that would not dock with the major connexin proteins endogenous to the mammalian brain. Though none of the protein candidates passed this level of screening, we believe several of these Cx34.7 and Cx35 mutants could potentially be used to enhance circuit editing approaches in *C. elegans.* Specifically, Cx34.7_K222Q_ and Cx35_E224H_, which failed to show homotypic docking, both docked with each other and with Cx36. On the other hand, Cx35_E224K_ and Cx35_E224R_, which did not show heterotypic docking with Cx36, both docked with Cx34.7_K222Q_. Thus, we believe that Cx34.7_K222Q_, Cx35_E224H_, Cx35_E224K_ and Cx35_E224R_ can be deployed combinatorially alongside Cx36 to facilitate multi-logic circuit editing in *C. elegans*.

We employed *in silico* modeling to address the unintended docking of our candidate proteins with Cx36, a feature that we believe would have impaired their application in mammals. Specifically, we developed a computational modeling pipeline to predict and simulate connexin hemichannel docking. We then employed this computational pipeline to probe the docking properties of the mutant proteins identified in our initial FETCH screen and to design new connexin mutants, *in silico.* At each step of our *in silico* design process, we validated the homotypic and heterotypic docking properties of our designed mutants by directly assaying them using our *in vitro* docking FETCH screen. Finally, we integrated all the mechanistic understanding of connexin docking interactions gained through this integrated approach to engineer Cx34.7_E214K/E223K_ and Cx35_K221E_, which show heterotypic, but not homotypic, docking *in vitro* and *in silico*. To our knowledge, this exclusively heterotypic docking profile has never been observed for pairs of connexin hemichannels. The docking specificity of Cx34.7_M1_ and Cx35_M1_ was validated *in vivo* in *C. elegans,* where we demonstrated that their heterotypic, but not homotypic expression in two distinct cells can edit the circuit between them and modulate behavior. We also confirmed the functionality of our mutant Cx34.7/Cx35 pair. This latter finding was exactly as we anticipated since the residues targeted for mutation were known to determine Cx hemichannel docking specificity, but not other physiological properties such as conductivity or gaiting, and both Cx34.7_WT_ and Cx35_WT_ successfully edited AFD/AIY. Critically, the circuit and behavioral outputs obtained using our final Cx34.7/Cx35 mutants were indistinguishable from those observed with Cx36/Cx36.

Since we also optimized the translational potential for LinCx by engineering Cx34.7 and Cx35 to disrupt their heterotypic hemichannel docking with other the major human connexin proteins Cx36 and Cx43, we believed that our tool can be broadly applicable to other preclinical model organisms including rodents and non-human primates. Indeed, we found strong sequence homology/identity of ELs across mammals (see supplemental Figure S2). To demonstrate this wide applicability, we validated the properties of LinCx directly in mice, where we found that LinCx successfully modified dynamics that reflect a microcircuit composed of two genetically distinct cell types. Specifically, deployment of Cx34_M1_ and Cx35_M1_ to prelimbic cortex pyramidal and parvalbumin-expressing interneurons, respectively, enhanced coupling between theta and high frequency oscillations in mice. This activity was modulated at the millisecond timescale. Strikingly, we also found that circuit editing was specific; theta and low gamma oscillation coupling, and power across the theta, gamma, and high frequency range were not impacted. Finally, we found that circuit editing was dependent on the orientation of the mutant hemichannels, providing evidence for rectification.

Within brain circuits, space is operationalized as the physical boundaries of individual cells, and time is operationalized as the sub-millisecond level electrical changes in those cells. Though many classic electrical stimulation approaches exhibit high temporal precision in their targeting, these techniques often stimulate volumes of brain tissue that include many brain cell-types and local axonal fibers of passage (see Supplemental Fig. S9; electrical stimulation). Preclinical approaches such as optogenetics provide a substantial improvement with regards to spatial targeting by enabling the selective stimulation of specific cell bodies (based on their genetic identities). Moreover, optogenetics modulates cells via temporally precise light pulses, maintaining the temporal precision of electrical stimulation. Nevertheless, both electrical and optogenetic stimulation bear substantial potential to override circuit computations, which integrate space and time across precise cell types, since these approaches are typically utilized under conditions that modulate the activity of many neurons concurrently and outside of the activity context of their inputs (see Supplemental Fig. S9; optogenetics - soma stimulation).

Over the last decade, several strategies have been employed across myriad studies to enhance the context precision of circuit targeting. One such strategy is based on selectively modulating projection neurons, whereby the inputs at a targeted brain site are selectively activated (Supplemental Fig. S9; optogenetics-projection stimulation) (Deisseroth, 2011). By directly modulating presynaptic nerve fibers, investigators can regulate the context (presynaptic neurotransmitter release) that drives the target cellular response. Nevertheless, this approach has the potential to profoundly alter physiological variables that define the context of the presynaptic neuron (and thus the circuit). For example, terminal stimulation can activate axonal collaterals of the presynaptic neuron thereby decreasing the spatial precision of circuit targeting, or it can induce retrograde activation of the presynaptic cell in a manner that disrupts the activation context of that neuron relative to its own inputs. This approach can also drive the activation of non-target circuits since inputs from a brain region can synapse onto multiple distinct cell types.

Other strategies utilized to enhance the context precision of circuit targeting include the stable step function opsins (SSFOs) (Yizhar et al., 2011) and designer receptors exclusively activated by designed drugs (DREADDs) (Armbruster et al., 2007), which function to increase the resting membrane potential of target cells. SSFOs and DREADDs maintain the cell type specific spatial precision characteristic of initial optogenetic targeting approaches. Moreover, because cells are rendered more likely to fire in response to their input signals under optimal conditions, these approaches provide improved temporal and context precision. Nevertheless, SSFOs and DREADDs can render target neurons more responsive to all their excitatory inputs (including those from non-targeted circuits), raising the potential of circuit-level off-target effects (see Supplemental Fig. S9; DREADDs). Thus, there is still demand for neuromodulation approaches that function within the spatial, temporal, and context constraints that together define brain circuit operation.

Another strategy to address limitations in context precision is to deliver stimulation within a closed loop framework. In this framework, neural activity is modulated based on the ongoing activity in the brain. For example, stimulating hippocampus at the peak vs. trough of the endogenous theta oscillatory cycle differentially impacts spatial memory (Siegle and Wilson, 2014). In our prior work exploring the circuitry underlying stress-induced behavioral adaptation, we stimulated neurons in medial dorsal thalamus while mice were engaged in a tail suspension test (Carlson *et al*., 2017). When we activated neurons based on ongoing oscillatory activity in infralimbic cortex during the test (i.e., closed loop circuit stimulation), mice showed decreased immobility. Conversely, when we stimulated these same cells using two distinct temporal patterns that were untimed to activity in infralimbic cortex, we observed either increased behavioral immobility, or no behavioral effect at all. Additionally, no behavioral effect was observed when we stimulated the terminals of infralimbic cortex neurons in medial dorsal thalamus. Together, these observations provided evidence that context precision serves as another critical axis of circuit operation in the brain that is orthogonal to both spatial and temporal precision. Moreover, these findings highlighted an ideal circuit and behavioral paradigm to test our new neuromodulation approach that was designed to preserve the spatial, temporal, and context precision of neural circuits.

Here, we utilized an integrated optogenetic stimulation and electrical recording approach to determine that LinCx expression potentiated the ILàMD circuit *in vivo*. We also found that this LinCx modulation of the ILà MD circuit decreased immobility during repeated tail suspension testing. Strikingly, this was a behavioral outcome that we could only previously achieve through closed loop stimulation. Interestingly, mice expressing LinCx exhibited normal behavior on the first day of tail suspension testing (t_24_=1.57; P=0.13 compared to non-edited mice using post-hoc testing with unpaired two-tailed t-test). This suggests that rather than suppressing acute stress behavior per se’, LinCx enhanced the compensatory function of endogenous ILàMD circuitry, ultimately suppressing behavioral adaptation in response to acute stress. As such, our findings suggest that LinCx likely enhances circuits within their physiological range, rather than driving them into supraphysiological states. This observation is supported by our prelimbic cortex micro-circuit editing findings which revealed that the majority of LinCx edited mice exhibited cross-frequency coupling that was less than the upper bound of the non-edited mice. Nevertheless, further testing is necessary to confirm this feature of LinCx.

Like established protein-based modulation tools such as optogenetics and DREADDs, LinCx can be targeted to precise cell types in mammals. However, LinCx builds upon these technologies by enabling each hemichannel to be expressed in a different cell type. The hemichannels expressed by these two distinct cell types then integrate *in vivo* to putatively form an electrical synapse. As such, LinCx offers unprecedented spatial precision compared to optogenetics and DREADDs in that it enables targeting of one of the specific spatial features that constrains circuits (e.g., the structural integration of two distinct cell types). LinCx is also designed to optimize the context precision of neuromodulation. Because this electrical synapse rectifies and it only forms between the target pre- and post-synaptic neuron, LinCx constrains the modulation of each individual post-synaptic neuron by the endogenous activity of its genetically defined pre-synaptic partner. This feature yields a level of temporal precision that mirrors the precision of endogenous brain activity, as confirmed by the enhanced millisecond-precision coupling of local neural activity to the phase of theta oscillations (i.e., periodicity of 100-250ms) observed in our LinCx edited mice. Critically, this feature also suggests that LinCx only potentiates circuit activity within the broader brain state context for which those circuits are typically engaged (i.e., the presynaptic neuron remains under the normal control of its own inputs). Finally, unlike established modulation approaches, LinCx does not require an exogenous actuator such as light, electricity, or an inert pharmacological compound. Rather, LinCx utilizes endogenous brain activity to modulate target neurons, yielding a tool for precise circuit editing.

The integrated engineering approach we utilized to develop LinCx (see Fig. 2E) can likely be deployed to develop a toolbox of connexin protein pairs that exhibit selective docking properties. Future work may also yield novel hemichannel pairs with customized conductance properties, mirroring approaches applied to modify the conductance of invertebrate electrical synapses (Shui et al., 2020). Thus, we believe that it may be possible to deploy multiple LinCx pairs in the same animal to simultaneously edit multiple circuits and ultimately regulate brain function. Critically, LinCx can also be deployed alongside other well established preclinical modulation approaches including DEADDS and optogenetics (as shown), enabling broad manipulation of brain networks across multiple scales of spatial, temporal, and context resolution concurrently.

### Limitations of the Study

There are several important limitations of our LinCx approach. First, like prior circuit editing approaches based on inserting gap junctions between specific cell types, our approach is only suitable for editing circuits composed of cells that make physical contact. Second, LinCx has the potential to yield mixed synapses (in neurons) for which chemical and electrical synapses operate in parallel. Since our connexin channels are chronically expressed, we anticipate that LinCx also induces changes in local chemical synapses. Indeed, this plasticity at chemical synapses may be fundamental to the physiological and behavioral changes we observed in our mouse assays.

We engineered Cx34.7 and Cx35 mutant proteins to be docking incompatible with Cx43 and Cx36. Since there are other connexin proteins expressed by mammals, we also used FETCH to screen our connexin mutants for heterotypic docking with other human connexin proteins. We observed FETCH scores for Cx31.3 and Cx37 that were higher than the docking incompatible pairs we used for our initial analysis (Supplemental Fig. S10), but lower than the docking compatible pairs. Cx31.3 is expressed in the parenchyma of the mammalian central nervous system (Lee et al., 2020), while Cx37 is not. Thus, future work is warranted to assess the docking compatibility of our mutant pairs with Cx31.3 in vivo, and the functional significance of any putative docking interactions, for CNS applications of LinCx.

Finally, we cannot exclude the possibility that our mutants oligomerize with endogenous Cx36 in mammals, yielding heteromeric hemichannels. Indeed, any such heteromeric channels may exhibit docking properties that are distinct from Cx36, Cx34.7_M1_, or Cx35_M1_ exclusive hemichannels (homomeric), ultimately limiting the functionality of LinCx across some neural circuits. Future work to assess and optimize the oligomerization specificity for Cx34.7_M1_ and Cx35_M1_ may further enhance the applicability and utility of LinCx to mammalian neural circuit editing.

## Conclusion

Direct stimulation of the brain is a well-established treatment for neurological and psychiatric disorders. For example, electrical convulsive therapy (ECT), which delivers energy to the whole brain, has remained the most effective treatment for major depressive disorder for nearly a century (Pagnin et al., 2004). Deep brain stimulation (DBS) of the sub thalamic nucleus is a widely utilized therapeutic for Parkinson’s disease (Liu et al., 2014). Nevertheless, both these modalities have important spatiotemporal constraints. ECT can induce short term cognitive dysfunction, likely due to the spatially untargeted nature of energy delivery to the brain, ultimately limiting its clinical use. On the other hand, DBS of the subthalamic nucleus has limited impact on the devastating emotional and cognitive symptoms that accompany Parkinson’s disease, likely due to the brain region targeting selectivity (Liu *et al*., 2014). While non-invasive techniques such as transcranial magnetic and focused ultrasound hold future promise for expanding the therapeutic landscape for brain disorders due to their increased clinical accessibility (these approaches do not require anesthesia/brain surgery), there continues to be demand for novel therapeutic approaches that modulate brain activity within both the spatial and temporal constrains of brain circuit activity.

We believe there is great potential for LinCx as a tool to probe the causal relationship between brain circuit function and behavior, and as a potential therapeutic approach to ameliorate human neuropsychiatric disorders. Moreover, there are many potential applications for LinCx technology that extend beyond the central nervous system. For example, future studies could optimize and deploy LinCx to the neuromuscular junction as a potentially treatment for Myasthenia Gravis, the cardiac nervous system as a potential treatment for arrhythmias, to the splenic nerve to regulate inflammation, the autonomic nervous system as a potential treatment for a range of disorders associated with gastrointestinal dysfunction, or to the integumentary system to ameliorate chronic pain.

## Materials and Methods

### Design of Cx34.7 and Cx35 mutant library

A semi-rational design approach was used to design the mutant library. Sequence alignments between the *Morone americana* connexins and the connexins for which the most structure-function data existed (Cx26, Cx32, Cx36, Cx40, and Cx43) were performed in ClustalW. Sites identified by previous studies as conferring specificity for docking were used as well as those identified by homology modeling from the structures of Cx26 (Koval *et al*., 2014). Specifically, we primarily focused on the extracellular loops and four residues at the interface in loop two, KEVE/KDVE (*M. americana* Cx34.7/Cx35) and one residue of E1. The homologous residues in other connexins had been demonstrated to be highly tolerant to mutation and critical for docking specificity (Jassim et al., 2016). Mutations were modeled in Swiss PDB Viewer using homology models of Cx34.7 and Cx35 from a Cx26 and Cx32 interface structure so as not to create mutations with obvious steric hindrance. A wide range of substitutions was made for these five residues of interest, including both those intended to introduce compatible electrostatic interactions as well as less likely candidates. Mutations were also created targeting other residues nearby and/or adjacent to these five for which there was some evidence in the literature that they contributed to docking specificity. However, our semi-rational approach was such that not as many variants were tried for these more distal site mutations and the mutations that were made in those sites were more conservative with regards to the steric and electrostatic properties of the change.

### Construct cloning and preparation

*Morone Americana* Cx34.7 and Cx35 cDNA constructs we initially acquired failed to efficiently express in in HEK 293FT cells. Thus, Connexin gene information was procured from the National Center for Biotechnology Information (NCBI, ncbi.nlm.nih.gov) and the Ensembl genome browser (ensembl.org). The human codon-optimized genes were ordered from Integrated DNA Technology as gBlocks Gene Fragments (IDT, idtdna.com). To generate constructs for transient transfection of HEK 293 FT cells, genes were subcloned into Emerald-N1 (addgene:53976) and piRFP670-N1 (addgene: 45457) vectors using In-Fusion cloning (takarabio.com), resulting in connexin fluorescent fusion proteins, specifically with the fluorescent proteins being adjoined to the connexin carboxy-terminus. Mutant constructs were generated by employing overlapping primers within standard Phusion polymerase PCR reactions to facilitate site-directed mutagenesis.

The Gateway recombination (Invitrogen) system was used to generate all Connexin 36, Cx34.7, Cx35, wild type and mutant protein *C. elegans* expression plasmids. For PCR-based cloning and subcloning of components into the Gateway system, either Phusion or Q5 High-Fidelity DNA-polymerase (NEB) was used for amplification. All components were sequenced within the respective Gateway entry vector prior to combining components into expression plasmids via a four-component Gateway system (Merritt and Seydoux, 2010). The different connexins versions were introduced into pDONR221a using a similar PCR-based strategy from plasmid sources (Chelur and Chalfie, 2007; Chen et al., 2013; Rabinowitch *et al*., 2014). Cell-specific promoters were introduced using the pENTR 50-TOPO vector (Invitrogen) after amplification from genomic DNA or precursor plasmids. Transgenic lines were created by microinjection into the distal gonad syncytium (Mello and Fire, 1995) and selected based on expression of one or more co-injection markers: Punc-122::GFP, Pelt-7::mCherry::NLS.

### Cell Culture

HEK 293FT cells were purchased from Thermo Fisher Scientific (cat# R70007) and were maintained according to manufacturer instructions. Briefly, cultures were grown in 10 cm tissue culture treated dishes in high-glucose DMEM (Sigma Aldrich, D5796) supplemented with 6mM L-glutamine, 0.1 mM MEM non-essential amino acids and 1 mM MEM sodium pyruvate in a 5% CO_2_, 37°C incubator. Cells were passaged via trypsinization every 2-3 days or until 60-80% confluency was reached.

### Transient Transfection

HEK 293FT cells were plated in 10ug/ml Fibronectin coated multi-well dishes to achieve ∼75% confluency after overnight incubation. For transfection, 250ng DNA was combined with polyethylenimine (PEI) diluted in Opti-MEM in a 1:3 ratio (µg of DNA: µL of PEI) and incubated at room temperature for 10 minutes. Following incubation, PEI-DNA complexes were added dropwise to wells of plated cells. Treated cells were then incubated at 37°C for 16-18 hrs, followed by media change. Expression of the connexin-FP constructs were evaluated at 24 hrs and 48 hrs post transfection via widefield or confocal microscopy and western blotting.

### Flow Enabled Tracking of Connexosomes in HEK 293FT cells (FETCH)

FETCH analysis is fundamentally a two-component system. To complete FETCH analysis, replica multi-well plates with HEK 293FT cells were transfected with either of the two components being evaluated. The media of transfected wells was changed 16-18 hrs post transfection and cells were trypsinized. Next, HEK 293FT cells expressing experimental connexin counterparts were combined. The entirety of combined samples was then plated onto new, 10ug/ml Fibronectin coated wells of the same size, resulting in hyperdensity and over confluency. Following co-plating, samples were incubated for ∼20-24 hrs, allowing cells to make contacts and potentially generate and internalize dually-labeled gap junctions. Samples were then trypsinized, resuspended in phosphate buffered saline with 10U/ml DNAse and fixed with paraformaldehyde (f/c of 1.5%). Co-plated samples in 96 well plates were resuspended to a final volume of ∼150 µL, samples from 24-well plates were resuspended to a final volume of ∼600 µL. Flow cytometry data was collected on either a BD FACSCanto II (2-color experiments and high-throughput 96-well plates; 488nm and 633nm lasers) which utilizes the BD FACSDiVa software. Samples were analyzed in two selection gates prior to their fluorescence evaluation. First, presumable HEK cells were identified by evaluating sample forward vs side scatter area. Next, single cells were selected by evaluating cells that maintained a linear correlation of forward scatter height to forward scatter area. Finally, the fluorescence profiles of each sample were generated.

### FETCH automated gating pipeline

Each FETCH experiment produces “*.fcs” files that contain all the channel data for fluorescence in the sample. Our automated pipeline loads these files, extracts FSC-A, SSC-A, and FSC-H. Depending on the machine used, we either load green channel as 1-A or as FITC-A. For the red channel, we can either have two (APC-A (RFP670) and PE-A (mApple)), or just one: 5-A(RFP670). Next, our code produces two matrices containing SSC-A with FSC-A, and FSC-A with FSC-H respectively.

Our first gate is drawn on the FSC-A vs SSC-A axes to exclude cellular debris which clusters in the lower left corner and the cells that are saturating the laser (at the max of both axes). On an FSC-A vs SSC-A plot, the cellular debris usually is smoothly transitioning into the population of intact cells, therefore we use a Gaussian kernel density estimator with the estimator bandwidth selection defined by the Scott’s Rule to draw contours around the data in SSC-A vs FSC-A matrix. We next use a set of heuristics to determine which of the contour lines should be used to define the first gate. Specifically, cellular debris usually clusters below 25000 on both axes, so any contour that includes values at or below is excluded. Similarly, any contour within a 1000 of the maximum value of each axis is also excluded. Of the remaining contours, the largest one is selected, and an oval equation is fitted to the points defining that contour to attenuate occasional protrusions of the contour that tap into cellular debris population in rare cases. The fitted oval becomes the first gate.

For all the elements inside of the first gate, a second gate is drawn in the axis of FSC-A and FSC-H to exclude non-single cells. For the second gate, first we fit a line to all the points. Next, for each point we find a norm to the fit line and find a standard deviation of all such norms. Using this standard deviation, we define a second gate 4 standard deviations away from the fitted line on both sides and exclude all the points outside of this gate.

Upon applying the first two gates, we finally plot the data with the red fluorophore on the y axis and the green fluorophore on the x axis. If a sample contains more than two fluorophores the last gate is drawn for each possible combination. Since some readings are below zero due to fluorescence compensation, we shift all the data points by the smallest value along both axes and then take a natural log of fluorescence levels. To achieve the optimal bandwidth for the kernel density estimation, we run a cross validation grid search algorithm on the points in the log space. Then we fit a gaussian kernel density with the bandwidth estimated to obtain density contours. For properly expressing samples, we expect a large population of untransfected cells in the bottom left quadrant of the plot, a population of cells strongly-expressing red fluorophore along the y axis, and a population of cells strongly expressing green fluorophore along the x axis. We expect autofluorescence to not exceed 500 on either axis, so the untransfected population is defined to be below this value along both axes. To draw a tighter bound of the untransfected population, we choose the first contour whose mean kernel density estimate (kde) value is at or above the 60th quantile (identified as a generalizable heuristic value) of the distribution of kde values within the largest contour which is at or below the autofluorescence cutoff. The top-most point of the tight contour defines the horizontal gate and the right-most point -- the vertical gate, separating the plot into four quadrants.

The upper left quadrant Q1 corresponds to the cells expressing just the red fluorophore, the upper right quadrant Q2 represents dual-colored cells, the lower right quadrant Q3 -- the cells expressing just the green fluorophore, and the lower left quadrant Q4 represents untransfected population. The FETCH score is defined as the portion of transfected cells that were dual-colored: Q2/(Q1 + Q2 + Q3).

Expecting approximately equal expression levels of each fluorophore, if the number of cells in Q1 is two or more times larger/smaller than the number of cells in Q3, the FETCH score is classified as “dubious” and marked accordingly in the output table. The “dubious” label is also given to samples that have less than 500 cells total after the application of the second gate and to the samples that failed at any of the steps in the pipeline (usually due to poor expression or the absence of cells in the sample). Code is available at: https://github.com/carlson-lab/FETCH.

### *In vitro* screening of Cx34.7 and Cx35 mutants for docking selectivity

For homotypic docking screening analysis, five FETCH replicates were obtained for each mutation. These scores were benchmarked against scores for Cx36 and Cx45 (FETCH=1.2±0.1%, N=54 replicates). For our heterotypic docking screening analysis, five replicates were obtained for each mutant pairs. These scores were then benchmarked against scores for wild type Cx34.7 and Cx35 (FETCH=14.7±0.4%, N=49 replicates).

To quantitatively determine whether a connexin pair docked, we determined FETCH scores for the dual fluorescence of cells under conditions where docking was not anticipated. These conditions included pairs of connexins previously established to not show docking: Cx36 and Cx45 (FETCH=0.7±0.0%, N=59 replicates), homotypic Cx23 (FETCH=0.9±0.4%, N=6), Cx36 and Cx43 (FETCH=1.2±0.2%, N=10), and under conditions for which cells were transfected with cytoplasmic fluorophores rather than tagged connexins (FETCH=4.4±0.6%, N=17). These 92 FETCH scores were used as the ‘known-negative’ distribution. FETCH scores from each experimental condition were then compared against the established negative score distribution using a one-tailed t-test, with a Bonferroni correction for the total number of experimental conditions tested. These FETCH replicates were independent of the replicates utilized for our screening analysis. Stats are reported as mean±s.e.m, and only uncorrected P values are reported through the text.

### Confocal Imaging Analysis of Gap Junction Partners

For imaging of putative gap junction partners, different populations of HEK 293FT cells were transfected with counterpart connexin proteins, incubated, and combined as described for FETCH analysis. Combined samples of HEK 293FT cells were co-plated onto 10 ug/ml Fibronectin coated 35 mm, glass-bottom Mattek dishes (cat# P35GC-1.5-14-C). Cells were imaged at ∼20 hrs post co-plating. Images were acquired on a Leica SP5 laser point scanning inverted confocal microscope using Argon/2, HeNe 594nm and HeNe633nm lasers, conventional fluorescence filters and a 63X, HCX PL APO W Corr CS, NA: 1.2, Water, DIC, WD: 0.22mm, objective. Images were taken with 1024 x 1024 pixel resolution at 200Hz frame rate.

For assessing Cx34.7_M1_::Cx35M1 expression *in vivo* in *C. elegans*, we imaged strain DCR8669 *olaEx5214 [Pgcy-8::CX34.7(E214K, E223K)::GFP; Pttx-3::CX35(K221E)::mCherry; Punc-122::GFP]*. L4 animals were mounted in 2% agarose in M9 buffer pads and anaesthetized with 10mM levamisol (Sigma). Confocal images were acquired with dual Hamamatsu ORCA-FUSIONBT SCMOS cameras on a Nikon Ti2-E Inverted Microscope using a confocal spinning disk CSU-W1 System, 488nm and 561nm laser lines and a CFI SR HP PLAN APO LAMBDA S 100xC SIL objective. Images were captured using the NIS-ELEMENTS software, with 2048px x 2048px, 16-bit depth, 300nm step size, 200ms of exposure time and enough sections to cover the whole worm depth.

### Protein modeling pipeline

Our protein modeling pipeline is based on previously published methodology (Lee, 2018) and integrates five components: 1) homology model generation, 2) embedding of proteins in a lipid bilayer and aqueous solution, 3) protein mutagenesis, 4) system minimization, equilibration, and molecular dynamics simulation production run, and 5) residue-wise energy calculation.

### Homology Modeling

We initially tested five homology modeling software suites: Robetta, SwissModel, Molecular Operating Environment (MOE; Chemical Computing Group ULC, Montreal, QC, Canada, H3A 2R7, 2021), I-Tasser, Phyre2 (Bertoni et al., 2017; Kallberg et al., 2012; Kelley et al., 2015; Roy et al., 2010; Song et al., 2013; Waterhouse et al., 2018; Yang et al., 2015; Yang and Zhang, 2015; Yang et al., 2011). A quality assessment suite, — MOLProbity (Chen et al., 2010; Davis et al., 2007; Williams et al., 2018) revealed that Robetta models outperformed the rest, based on a set of standard metrics (Ramachandran plot outliers, clashscore, poor rotamers, bad bonds/angles, etc).

Since our aim was to model the extracellular loops responsible for connexin hemichannel docking, we picked all the resolved connexin structures that possessed a high degree of extracellular loop homology to our connexins of interest as the inputs for Robetta. The top homolog hits were generally the same for the three Cxs of interest: Connexin-26 Bound to Calcium (5er7.1) (Bennett *et al*., 2016), Human Connexin-26 (Calcium-free) (5era) (Bennett *et al*., 2016), Structure of connexin-46 intercellular gap junction channel at 3.4 angstrom resolution by cryoEM (6mhq) (Myers *et al*., 2018), Structure of connexin-50 intercellular gap junction channel at 3.4 angstrom resolution by cryoEM (6mhy) (Myers *et al*., 2018), Structure of the connexin-26 gap junction channel at 3.5 angstrom resolution (2zw3) (Maeda *et al*., 2009). Cx34.7 and Cx35 wild type sequences had the greatest homology degree with 6mhq, while Cx36 was most homologous to 5er7.1. We generated three wild type hemichannels for Cx34.7, Cx35, and Cx36.

### System Assembly

Next, we assembled hemichannels into homotypic and heterotypic gap junctions, embedded them in two double bilayers, dissolved them in water, and added appropriate ion concentrations for the extracellular and two intracellular compartments. The primary software suite used for this modeling step was VMD (Eargle et al., 2006; Humphrey et al., 1996). We also utilized CHARMM GUI to generate the naturalistic model of a region of a double bilayer (Jo et al., 2007; Jo et al., 2008; Jo et al., 2009; Lee et al., 2019; Wu et al., 2014). Membrane components were then selected in appropriate proportions to resemble experimentally-derived data from a neuronal axonal membrane.

Specifically, since Robetta was unable to model the full gap junction, we merged hemichannels into full homotypic/heterotypic gap junctions in a semi-automated way. First, to make homotypic gap junctions, we loaded the two-homology models for a hemichannel. We then aligned them using the center of mass of the extracellular loops. A slight rotation along the z axis was implemented for several pairs to optimize their fit. To make heterotypic gap junctions, we created a homotypic gap junction for each hemichannel, aligned the extracellular loops for the two homotypic gap junctions, then removed an opposing hemichannel from each homotypic gap junction (leaving the two different hemichannels aligned). Next, using the constructed gap junction, we aligned two pre-made membrane bilayers with the center of mass assigned as each embedded hemichannel. We then removed membrane molecules that overlapped with the hemichannel or the hemichannel pore. Next, we dissolved the system in water, and removed water that overlapped with the lipid bilayer. Extracellular water was then separated to a new file, where Na+, K+, Cl-, and Ca2+ ions were added to yield concentrations mirroring the extracellular environment of mammalian neurons (Alberts, 2015). Finally, Na+, K+, Cl-, and Ca2+ ions were added to the intracellular space to mirror the intracellular environment of mammalian neurons, and the files containing the embedded connexin hemichannels and extracellular water were merged. Notably, these stages were automated yielding a streamlined progression from a protein-only hemichannel model to a fully embedded gap junction model ready for subsequent simulation and/or mutagenesis.

### Mutagenesis

We developed a python command-line tool that utilizes VMD to generate mutation configuration files for subsequent MD simulation. Here, we simply specified the connexin hemichannels of interest and the position at which a specific mutation should be introduced.

### System Minimization, Equilibration, and MD Simulation Production Run

Next, we minimized atomic energies, equilibrated the system, and ran the stable system in a production simulation run. Specifically, MD simulation was performed using NAMD *(Phillips et al., 2005)* and was divided into five steps:

1. *Melt lipid tails while keeping remaining atoms fixed (simulate for 0.5 ns)*
2. *Minimize the system, then allow the bilayers and solutions to take natural conformation while keeping gap junction fixed (split in two stages to accommodate reduction in volume of relaxing system; simulate 0.5 ns total)*
3. *Release the gap junction and equilibrate the whole system (simulate 0.5 ns)*
4. *Run minimized and equilibrated system in a production run (simulate 0.5 ns)*

Though MD simulation (step 4) is highly reliant on the input file provided by the *System Assembly* process, these steps render the simulation much more robust to modeling imperfections. For example, the membrane model developed though System Assembly is very rigid and has the potential to behave like a solid rather than like a liquid. Thus, melting the lipid tails encourages the model to embody a liquid. Similarly, many atoms in the input file may have unnatural initial energies, such that if they are all released at once, they would start moving at high velocities and the simulation would fail. Therefore, bringing the system to a local energy minimum increases stability. Removing constraints on the water and lipids enables them to surround the gap junction in a naturalistic form. Finally, releasing the constraints on the gap junction enables it to take the most energetically stable conformation given the environment.

### Energy Calculation

To predict the residues that play a prominent role in docking, we quantified all non-bonding interactions between the two connexin hemichannels at key residues on the extracellular loops. Output from the MD simulation was loaded into the VMD “NAMD Energy” plugin. We then calculated nonbonding energies for all residues on each hemichannel that were within 12 angstroms of at least one residue on the other hemichannel. For each residue pair we then averaged energies across the 250 simulation frames

### *C. elegans* strains and genetics

Nematodes were maintained at 20°C on NGM plates seeded with a lawn of *Escherichia coli* strain OP50 using standard methods (Brenner, 1974). All worm experiments were performed using one-day-old adult hermaphrodites. The strains used in this study are listed in the Strain Table (supplemental table S2). All thermotaxis behavioral assays, calcium imaging experiments and generation of transgenic lines were performed as previously described in Hawk et al. 2018 (Hawk *et al*., 2018), with minor modifications as outlined in the following subsections.

### Thermotaxis Behavioral Assay

Animals were grown and assayed as previously described in Hawk et al. 2018 (Hawk *et al*., 2018; Luo et al., 2014). Briefly, after being reared at 20°C, animals were trained at 15°C for 4 hours prior to testing. Their migration tracks were analyzed as previously described (Gershow et al., 2012; Hawk *et al*., 2018; Luo *et al*., 2014). Each behavioral arena was split in half along the temperature gradient using a thin and clear plastic divider. This allowed for wild type controls to be assayed on one half of the arena and connexin-expressing animals on the other half, simultaneously.

### Calcium Imaging in *C. elegans*

Imaging calcium dynamics for assaying AFD-AIY functional coupling was performed as previously described (Hawk *et al*., 2018), with the following modifications: a Leica DM6B was used instead of Leica DM5500, and image acquisition was performed using MicroManager (Edelstein et al., 2014). Segmentation into regions of interest and downstream data processing was performed using FIJI (Schindelin et al., 2012) and custom scripts written in MATLAB (MathWorks) were used as detailed previously (Hawk *et al*., 2018). For analyses of AFD calcium transients, we generated and measured an ROI around a single AFD soma per animal. For analyses of AIY calcium dynamics, we generated and quantified an ROI at the synaptic subcellular region known as Zone 2 (Colon-Ramos et al., 2007). Responses were scored as the initial rise of the AFD or AIY calcium signal as determined by a blind human observer. The genetic background for the AFD and some of the AIY calcium imaging lines used in this study (control and experimental) contained *olaIs23*, a caPKC-1 GOF mutation. This was done to match prior work (Hawk *et al*., 2018) in which Connexin 36 was demonstrated to evoke AFD-locked responses in AIY compared to caPKC-1 animals without Connexin 36.

### Vertebrate Animal Care and Use

Male B6.129P2-*Pvalb^tm1(cre)Arbr^*/J (PV-Cre mice) and female C57BL/6J mice (Stock No: 017320 and 000664, respectively) purchased from Jackson labs were bred to generate the male and female (N=28 total virally injected and implanted mice) PV-Cre heterozygous mice subjected to the prefrontal cortex PYR↔ PV+ editing experiment. Male C57BL/6J mice (N=29) purchased from Jackson were utilized for non-edited controls. Mice were housed three-five/cage on a 12-hour light/dark cycle and maintained in a humidity- and temperature-controlled room with water and food available ad libitum. Neural recordings were conducted during the dark cycle (Zeitgeber time: 13-19), given prior evidence that electrical synapse conductance can be diminished in the retina via circadian regulation (Zhang et al., 2015). Inbred BALB/cJ male mice (strain: 000651) purchased from the Jackson Labs were used for infralimbic cortex à medial thalamus circuit editing experiments. We chose this strain and sex of mice to mirror our previous study which optogenetically targeted the infralimbic cortex à medial thalamus circuit (Carlson *et al*., 2017). Behavioral and physiological experiments were conducted during the dark cycle (Zeitgeber time: 13-22). All vertebrate animal studies were conducted with approved protocols from the Duke University Institutional Animal Care and Use Committees and were in accordance with the NIH guidelines for the Care and Use of Laboratory Animals.

### Mouse viral injection surgeries

For Cx35WT experiment, C57BL/6J mice were in anesthetized with isoflurane (1%), placed in a stereotaxic device, and injected with AAV9-CaMKII-Cx35_M1_-mEmerald (titer: 6.9×10^12^ vg/ml) or AAV9-CaMKII-Cx35_WT_-mApple (titer: 9.3 x 10^12 vg/ml), based on stereotaxic coordinates measured from bregma at the skull to target ventral hippocampus (−3.3mm, 3.0mm ML, −3.5mm DV from the dura for male mice, and −3.3mm, 3.0mm ML, −3.5mm DV from the dura for female mice). A total of 0.3µL or 0.5 µL viral solution was delivered to each hemisphere over 10 minutes using a 5µL Hamilton syringe.

For PYR-PV+ editing experiment, PV-Cre mice were anesthetized with isoflurane (1%), placed in a stereotaxic device, and injected with a 1:1 solution of AAV9-CaMKII-Cx34.7_M1_-mEmerald (titer: 5.0×10^12^ vg/ml) and AAV9-Ef1α-DIO-Cx35_M1_-mApple (titer: 1.3×10^13^ vg/ml), based on stereotaxic coordinates measured from bregma at the skull to target prelimbic cortex bilaterally (PrL: 2.1mm AP, 0.65mm ML, −1.45mm DV from the dura at a 21⁰ angle for male mice, and PrL: 2.05mm AP, 0.62mm ML, −1.41mm DV from the dura at a 21⁰ angle for female mice). A total of 1µL viral solution was delivered to each hemisphere over 10 minutes using a 5µL Hamilton syringe. This strategy yielded expression of Cx34.7_M1_ by cortical excitatory and inhibitory neurons, and expression of Cx35_M1_ solely by PV+ expressing interneurons. Control mice were injected with a 1:1 solution of AAV9-CaMKII-Cx34.7_M1_-mEmerald and AAV9-Ef1α-DIO-Cx34.7_M1_-mApple (titer: 1.1×10^13^ vg/ml) or a 1:1 solution of AAV9-CaMKII-Cx35_M1_-mEmerald (titer: 6.9×10^12^ vg/ml) and AAV9-Ef1α-DIO-Cx35_M1_-mApple. These viruses were created in the Duke Viral Vector Core. Viral injections were performed in male and female mice at age 2.5-5 months, and viral manipulations were balanced across cages and sex.

For IL->MD behavioral experiment, BALB/cJ mice were anesthetized with isoflurane (1%), placed in a stereotaxic device, and injected with AAV9-CaMKII-Cx34.7_M1_-mEmerald (titer: 5.0×10^12^ vg/ml) based on stereotaxic coordinates measured from bregma at the skull to target infralimbic cortex bilaterally (IL: 1.7mm AP, 0.72mm ML, −2.03mm DV from the dura at an angle of 10°). A total of 0.5µL viral solution was delivered to each hemisphere over 5 minutes using a 5µL Hamilton syringe which was left in place for an additional 10 minutes prior to removal. Three weeks later (see supplemental Figure S4), mice were injected with AAV9-CaMKII-Cx35_M1_-mApple (titer: 6.9×10^12^ vg/ml) based on stereotaxic coordinates measured from bregma at the skull to target medial dorsal thalamus bilaterally (MD: −1.58mm AP, 0.5mm ML, −2.88mm DV from the dura at an angle of 10°). Control mice were injected with AAV9-CaMKII-Cx34.7_M1_-mEmerald in both IL and MD, or AAV9-CaMKII-Cx35_M1_ in both IL and MD, to express the synthetic hemichannels in non-docking homotypic configurations. Viral injections were performed in male mice at age 3 months. For the ILàMD circuit interrogation study, mice were injected with a 1:1 solution of AAV9-CaMKII-Cx34.7_M1_-mEmerald (titer: 2.3×10^13^ vg/ml) and AAV9-CamKII-ChR2-EYFP (titer: ≥ 1×10¹³vg/ml) to target IL unilaterally (1.7mm AP, −0.72mm ML, measured from bregma; −2.03mm DV from the dura at an angle of 10°). A total of 0.5µL viral solution was delivered at a rate of 100nL/minute. Three weeks after the first surgery, mice were again anesthetized, placed in a stereotaxic device and injected with either AAV9-CaMKII-Cx35_M1_-mApple (titer:3.16×10^13 vg/ml) or AAV9-CaMKII-EGFP (titer: 2.3×10^13^ vg/ml) to target medial dorsal thalamus (MD) unilaterally. A total of 0.5µL viral solution was delivered. Viruses were infused at a rate of 100nL/minute.

### Electrode Implantation Surgery

For PYR-PV+ experiment, PV-Cre mice mice were anesthetized with isoflurane (1.0%), placed in a stereotaxic device, and metal ground screws were secured to the cranium. A total of 8 tungsten microwires were implanted in prelimbic cortex (centered at 1.8mm AP, ±0.25mm ML, −1.75mm DV from the dura for male mice; centered at 1.76mm AP, ±0.25mm ML, −1.71mm DV from the dura for female mice). C57BL/6J control mice were implanted at 2 months. A total of 32 tungsten microwires were arranged in our previously described multi-limbic circuit recording design(Mague et al., 2020). Briefly, bundles were implanted to target basolateral and central amygdala (AMY), medial dorsal thalamus (MD), nucleus accumbens core and shell (NAc), VTA, medial prefrontal cortex (mPFC), and VHip were centered based on stereotaxic coordinates measured from bregma (Amy: −1.4mm AP, 2.9 mm ML, −3.85 mm DV from the dura; MD: −1.58mm AP, 0.3 mm ML, −2.88 mm DV from the dura; VTA: −3.5mm AP, ±0.25 mm ML, −4.25 mm DV from the dura; VHip: −3.3mm AP, 3.0mm ML, −3.75mm DV from the dura; mPFC: 1.62mm AP, ±0.25mm ML, 2.25mm DV from the dura; NAc: 1.3mm AP, 2.25mm ML, −4.1 mm DV from the dura, implanted at an angle of 22.1°). We targeted cingulate cortex, prelimbic cortex, infralimbic cortex using the mPFC bundle by building a 0.5mm and 1.1mm DV stagger into our electrode bundle microwires. Animals were implanted bilaterally in mPFC and VTA. All other bundles were implanted in the left hemisphere. The NAc bundle included a 0.6mm DV stagger such that wires were distributed across NAc core and shell. We targeted BLA and CeA by building a 0.5mm ML stagger and 0.3mm DV stagger into our AMY electrode bundle(Mague *et al*., 2020).

For ILàMD circuit interrogation study, 16 tungsten microwires were arranged into two bundles to target IL (8 wires) and MD (8 wires). The IL bundle was also built with an optical fiber (MFC_100/125-0.22_8.0mm_MF2.5_FLT, Doric Lenses) 0.5mm above the tip of the wires as previously described (Carlson et al, Biological Psychiatry). An optic fiber was also built into the MD microwire bundle for two-thirds of animals, though not utilized for this study. Mice were anesthetized as described above, and metal ground screws were secured to the cranium. Bundles were implanted in IL (1.7mm AP, −0.15mm ML, from bregma; −2.25mm DV from the dura) and MD (centered at 1.58mm AP, −0.5mm ML, −2.88mm DV from the dura).

### Data acquisition for PrL microcircuit editing

Neural recordings experiments were performed in PV-Cre mice one week after implantation surgery and blind to viral group. Mice were habituated to the recording room for at least 60 minutes prior to testing. PV-Cre mice were connected to a headstage (Blackrock Microsystems, UT, USA) without anesthesia, and given a single saline injection (10mL/kg mouse, intraperitoneally). Notably, these saline injections were performed to facilitate comparison of acquired neural data with future drug studies. Twenty-five minutes later, mice were placed in a 17.5in × 17.5in × 11.75in (L×W×H) chamber for 60 minutes. Recordings were conducted under low illumination conditions (1-2 lux), and only data from the first 10 minutes of exposure to the open field were used for neurophysiological analysis. For C57BL/6J control mice, experiments were performed at least two weeks following implantation surgery. Mice were habituated to the recording room for at least 60 minutes prior to testing, and headstages were connected without anesthesia. Twenty-six male mice were placed in a 17.5in × 17.5in × 11.75in (L×W×H) chamber for 10 minutes, and three mice were recorded in a 19.5in × 12in (D×H) circular chamber. Six of these mice were injected with (10mL/kg mouse, intraperitoneally) 30 minutes prior to recordings, and all recordings were conducted under an illumination of 125 lux.

Neuronal activity was sampled at 30kHz using the Cerebus acquisition system (Blackrock Microsystems Inc., UT). Local field potentials (LFPs) were bandpass filtered at 0.5–250Hz and stored at 1000Hz. All neurophysiological recordings were referenced to a ground wire connected to both ground screws, and an online noise cancellation algorithm was applied to reduce 60Hz artifact.

### Determination of LFP cross frequency phase coupling and spectral power

Signals recorded from all viable implanted microwires were used for analysis. Local field potentials were filtered using 4th order Butterworth bandpass filters designed to isolate theta (4-10Hz) prefrontal cortex oscillations and high frequency oscillations (80-200Hz). The instantaneous amplitude and phase of the filtered LFPs were then determined using the Hilbert transform, and the Modulation index was calculated for each LFP channel using the MATLAB (The MathWorks, Inc., Natick, MA) code provided by Canolty et al (Canolty *et al*., 2006). Briefly, a continuous variable z(t) is defined as a function of the instantaneous theta phase and instantaneous gamma amplitude such that z(t) = A_G_(t)*e^iϕ^_TH_^(t)^, where A_G_ is the instantaneous gamma oscillatory amplitude, e^iϕ^_TH_ is a function of the instantaneous theta oscillatory phase. A time lag is then introduced between the instantaneous amplitude and theta phase values such that z_surr_ is parameterized by both time and the offset between the two variables, z_surr_ = A_HG_(t+τ)*e^iϕ^_TH_^(t)^. The modulus of the first moment of z(t), compared to the distribution of surrogate lengths, provides a measure of coupling strength. The normalized z-scored length, or Modulation index, is then defined as M_NORM_ = (M_RAW_-µ)/σ, where M_RAW_ is the modulus of the first moment of z(t), µ is the mean of the surrogate lengths, and σ is their standard deviation (Canolty *et al*., 2006; Carlson *et al*., 2017; Dzirasa *et al*., 2010). The modulation index scores were averaged across all implanted channels for each mouse (∼7.3 channels for each PV-Cre mouse, and 2 channels implanted bilaterally for each C57BL/BJ control mouse), yielding a single score per animal.

To quantify LFP oscillatory power, a sliding window Fourier transform with Hamming window was applied to the LFP signal using MATLAB. Data was analyzed with a 1 second window, 1 second step, and a frequency resolution of 1Hz. Signals were averaged across time windows and frequencies used for CFPC analysis, and then across all microwires, yielding a signal measure per animal.

### Data acquisition and analysis for *in vivo* IL-MD circuit interrogation

Experiments were conducted during two sessions: three weeks and five days after the first viral surgery, and again five weeks after the first viral surgery. A 473nm laser (CrystaLaser LC, DL473-025-O, CL-2005 Laser Power Supply) was calibrated using an optical power meter (ThorLabs PM100D) to deliver output at 3mW, 1mW, 0.75mW, 0.5mW, and 0.25mW. Laser stimulation was triggered using analog output from the Cerebus System (Blackrock Microsystems, UT, USA). Prior to recordings, mice were connected to a headstage and optic fiber without anesthesia and placed in a new cage.

Local field potential activity was acquired, as described above, concurrently with an analog signal corresponding to the laser TTL pulse. After a 10-minute baseline recording period, mice received repeated 10ms pulses of light in IL. Pulses were delivered with pseudorandomized inter-stimulus intervals ranging between 10 and 24 seconds. During the first session, mice were stimulated with light intensities of 0.25, 0.5, 0.75, and 1mW (30 trials each). We then determined the portion of mice that showed clear evoked responses at each light threshold. Light intensities were presented in pseudorandomized order. Stimulation protocol was fully automated using a Matlab script which will be available from https://github.com/carlson-lab/OptoLinCx.

Since our objective was to determine the impact of LinCx expression on ILàMD circuit physiology, our experimental approach required an IL stimulation intensity that was strong enough to evoke a potential in MD but did not saturate the elicited MD response. When we analyzed data from the first session (as described below), we observed that most animals failed to show an evoked potential in MD that was greater than 50µV using 0.25, 0.5, or 0.75mW stimulation. Thus, we utilized the 1mW IL stimulation (where 13/21 mice showed mean evoked responses to >50mW; see Supplemental Figure S7), to directly assess the impact of LinCx expression of ILàMD circuit function. We then added 30 trials of 3mW IL stimulation to the second session. Here, we reasoned that mice which failed to show clear evoked responses in MD in responses to 3mW stimulation (>75µW) would be below the detection threshold for determining the impact of LinCx expression at 1mW stimulation. Note that these experimental design criteria were implemented in an initial cohort of mice prior to histological confirmation of viral expression and electrode placement. Identical experimental parameters were then utilized for a second cohort of mice.

Custom Python scripts were used to analyze raw recordings (https://github.com/carlson-lab/OptoLinCx). First, the code loads raw NSX files and filters the local field potentials with a forward and backward second-order IIR notch filter at 60Hz, with a quality factor of 30, and a Butterworth analog highpass filter of order 5, with a cutoff of 15 Hz using Scipy implementation. After filtering, the average voltage in the −200ms to 1ms pre-stimulus window was subtracted out, normalizing each evoked response to 0mV. Trials were excluded if a given trace had more than 50% of the time points identified as outliers based on each point being assessed to be within 1.5 interquartile range (IQR) between the 25th and 75th quartile. The remaining trials were averaged within each channel and for each stimulus intensity. Channels that showed a pre-stimulus dominant frequency with peaks larger than 25μV were removed from subsequent analysis. Only animals with at least fifteen 1mW trials acquired during both sessions were utilized. Three mice that failed to show a robust response in MD to the 3mW stimulus were removed from subsequent analysis.

The mean of the peak amplitude in the 0 to 10ms window, averaged across all IL microwires, and in the 10ms to 25ms window, averaged across all MD microwires, was used for comparisons across groups. We reasoned that if an increase in the amplitude of MD evoked potentials was solely driven by an increase in IL evoked activity, we would observe a direct correlation between the two variables across animals. We found no such relationship between the amplitude change of MD and IL evoked potentials across the mice that expressed Cx35_M1_ (P=0.70 using Pearson correlation). Thus, our analysis was performed independently for each brain regions.

### Behavioral testing of ILàMD circuit manipulation

All behavioral testing was conducted under low illumination conditions (1-2 lux). Two weeks after the second viral surgery, mice were initially placed in a 17.5in × 17.5in × 11.75in (L×W×H) chamber for five minutes of open field testing. Mice were then suspended 1cm from the tip of their tail for six minutes for the tail suspension assay. Open field and tail suspension behavioral data were acquired during a single testing session, and the behavior testing session was repeated the next day. Testing sessions were video recorded, and open field and tail suspension behavior was analyzed using Ethovision XT 12 (Noldus, Wageniingen, the Netherlands). Behavioral experiments and subsequent video analyses were performed blind to group.

### Histology for mouse studies

Mice were perfused transcardially with ice-cold PBS followed by 4% PFA in PBS (EM Sciences, Hatfield, PA). The brains were harvested and coronally sliced in 1X PBS at 35µm using a vibratome (Vibratome Series 3000 Plus, The Vibratome Company, St. Louis, MO) and mounted onto positively charged slides using a mild acetate buffer (82.4mM Sodium Acetate, 17.6mM acetic acid) or 1X PBS solution. Brain slices were covered with DAPI-Mowiol mounting solution [Glycerol, puriss. p.a., Mowiol 4-88 (Sigma-Aldrich, St. Louis, MO), and 0.2M Tris-Cl pH 8.5, DAPI (Sigma-Aldrich, St. Louis, MO)] and cover slipped prior to imaging. Images were obtained using a Nikon Eclipse fluorescence microscope at 4x magnification with illumination source power and exposure kept consistent between samples. For the PrL PYRàPV+ study, we only used mice with confirmed bilateral connexin expression for neurophysiological analysis(N=26/29).

We also performed immunohistochemistry to assess the specificity of our viral targeting approach for the PrL PYRàPV+. Briefly, we injected three C57BL6/J mice in PrL with AAV-CamKII-Cx34.7_M1_-mEmerald. Following a 4-week expression period, brains were removed, sliced, and stained. We performed immunohistochemistry using anti-PV antibody and AlexaFluor 568 dye. Neurons were identified by DAPI expression and a diameter > 13µm. Green fluorescence (indicating viral expression) and red fluorescence (indicating PV) was then quantified across all neurons in PrL. Using this approach, we found that 4.3% of all cells that expressed Cx34.7_M1_ also expressed PV+, and 15.5% of PV+ neurons expressed Cx35_M1_. Thus, most of the cells that expressed Cx34.7_M1_ were excitatory. Nevertheless, AAV-CaMKII was not selective for excitatory neurons as previously reported. Indeed, we observed substantial labelling of PV neurons near the injection site.

For the ILàMD behavioral study, we used mice that showed bilateral connexin expression in one region, and at least unilateral expression in the other region (N=26/35). We chose this strategy since our prior work indicated that unilateral optogenetic stimulation of the ILàMD circuit was sufficient to alter stress related behavior in the tail suspension assay. For the ILàMD behavioral study, histology was performed as noted above to confirm electrode placement, EYFP/mEmerald trafficking to MD, and mApple expression in MD. In a subset of animals, verification of MD expression required additional staining against mApple. Here, we used Rabbit primary antibody against RFP (Rockland, 600-401-379) and secondary antibody with Alexa Fluor 568 (Abcam, ab175471). ChR2 expression was confirmed via electrophysiology (neural responses to blue light). Only mice with accurate targeting of both viral sites and electrodes were used for analysis.

## Acknowledgements

The authors would like to acknowledge Dr. Terry Oas, Dr. Mike Tadross, Dr. Nicole Calakos, Dr. Marc Caron, Dr. David Carlson, and Dr. Fan Wang for helpful discussion and/or guidance; Dr. Kayla Lemons, Dr. Gary Sutherland, and Reagan Portelance for technical assistance; and Dr. John O’Brien for supplying reagents. The authors would like to acknowledge the generous funding that has supported this work: Ernest E. Just Life Science Institute Postdoctoral Research Fellowship; Burroughs Wellcome Fund Postdoctoral Enrichment Program Grant (E.R.); Hartwell Postdoctoral Research Fellowship (E.R.); SOM Core Facility Voucher Program (E.R. and K.D.); NIH grants R01NS076558 and DP1NS111778 (D.C.R.); HHMI Scholar Award (D.C.R.); NIH grants U01HL134764, R01HL132389, and R01HL126524 (N.B.); Duke University School of Medicine MedX Grant (K.D. and N.B.); Duke University Chancellor’s Discovery award (K.D. and N.B.); Duke University School of Medicine Kahn Neurotechnology Grant (E.R., N.B., and K.D.); NIH BRAIN Initiative Grant R21EY029451 (K.D. and N.B.); Hope for Depression Research Grant, and NIH Grants R01MH120158, R01MH125430 and 1DP1MH132709 (K.D.). A special thanks to Freeman A. Hrabowski, Robert and Jane Meyerhoff, and the Meyerhoff Scholarship Program.

## Author contributions

Conceptualization and Methodology – E.R., K.C., E.W., A.A.P., D.C.R., R.H., and K.D.; Formal Analysis – E.R., K.C., G.E.T., E.W., T.R., K.K.C.W., D.H., A.A.P., S.D.M., and D.C.R., K.D.; Investigation – E.R., K.C., G.E.T., E.W., R.B., T.R., E.A., K.K.C.W., D.H., H.S., S.D.M., R.H., A.A.P.; Resources – E.R., N.B., D.C.R., K.D.; Writing – Original Draft, E.R., K.C., E.W., A.A.P., D.C.R., N.B., K.D. Writing – Review & Editing, E.R., K.C., G.E.T., E.W., T.R., A.A.P., K.K.C.W., D.C.R., S.D.M., R.H., N.B, K.D. Visualization – E.R., K.C., G.E.T., E.W., T.R., A.A.P., D.C.R., K.D.; Supervision – S.D.M., D.C.R., N.B. and K.D.; Project Administration and Funding Acquisition – E.R., D.C.R., R.H., N.B., and K.D. See Supplemental materials for detailed author contributions.

