## Supplemental Materials for "Long-term precision editing of neural circuits in mammals using engineered gap junction hemichannels"

**Supplementary Table S1, Detailed Author Contributions**

| | |
|---|---|
| Elizabeth Ransey | Conceived FETCH methodology as approach for screening connexin mutant docking; coordinated and performed connexin DNA construct cloning; performed all FETCH experiments and subsequent microscopy; conceived strategy to employ computational models for rational design; coordinated design of computationally guided mutations; designed and cloned AAV vector plasmids; secured funding and resources; prepared figures; wrote original draft of the introduction, *in vitro* study results and associated materials and methods, and discussion; and revised the paper. |
| Kirill Chesnov | Conceived and developed the computational modeling pipeline to enable rational design; performed all computational modeling analysis, conceived methodology for quantification of FETCH results, analyzed FETCH results; discovered motif interaction schema that guided Cx34.7, Cx35, and Cx36 docking; Designed experiment testing wild-type Cx35 *in vivo* in mice with KD, performed and analyzed data from wild-type Cx35 *in vivo* experiment; Designed and performed circuit interrogation study using optogenetics with KD, performed all analysis for optogenetic circuit interrogation study, coordinated and performed histological analysis for all mouse editing experiments, performed detailed immunohistological quantification for PYR-PV+ experiment; prepared figures; wrote original draft of *in silico* study *results* and associated materials and methods, and discussion; and revised the paper. |
| Gwenaëlle Thomas | Initiated research on gap junction structural/functional properties across animal species, including target residues to alter docking properties; systematically organized findings, selected candidate proteins as substrates for LinCx, and presented recommendations (to RH and KD) with EA, RB, and HS; performed Cx34.7 expression and trafficking study in mice (viral surgeries and histological analysis); performed viral surgeries for PYR→PV+ mouse editing studies; collected and analyzed neurophysiological data for PYR→PV+ editing study with S.D.M. and K.K.W., performed viral surgeries and collected data for infralimbic cortex→medial dorsal thalamus editing behavioral experiments with KKW; analyzed data for infralimbic cortex→medial dorsal thalamus behavioral experiment; prepared figures. |
| Elias Wisdom | Conceived *C elegans* neurophysiological and behavioral experiments with AAP; Generated Cx34.7 and Cx35 wild-type and mutant expression vectors for *C. elegans*; performed *C. elegans* experiments testing wild-type and mutant Cx34.7 and Cx35 under homotypic and heterotypic expression conditions and analyzed results; prepared figures; wrote original draft of *in vivo* study results and associated methods with A.A.P. and D.C.; and revised the paper. |
| Agustin Almoril-Porras | Conceived *C elegans* neurophysiological and behavioral experiments with EW; generated *C elegans strains*; generated Cx36 and Cx43 expression vectors for *C. elegans;* performed *C. elegans* experiments testing mutant Cx34.7 and Cx35 against Cx36 and Cx43. under heterotypic expression conditions and analyzed results; assisted in calcium imaging analyses; coordinated all C. elegans experiments; prepared figures; wrote original draft of *in vivo* study results and associated methods with E.W. and D.C.R.; and revised the paper. |
| Ryan Bowman | Researched gap junction structural/functional properties across animal species, systematically organized findings, selected candidate proteins as substrates for |

| | |
|---|---|
| | LinCx, and presented recommendations (to RH and KD) with EA, GET, and HS; Cloned connexin DNA constructs; Tested in vitro methods for evaluating gap function. |
| Tatiana Rodriguez | Analyzed FETCH results; Cloned connexin DNA constructs; prepared figures; revised the paper. |
| Elise Adamson | Researched gap junction structural/functional properties across animal species, systematically organized findings, selected candidate proteins as substrates for LinCx, and presented recommendations (to RH and KD) with GET, HS, and RB; Cloned connexin DNA constructs; |
| Kathryn Katsue Walder-Christensen | Planned PYR→PV+ and IL→MD (behavioral) experiments with G.E.T., S.D.M., K.D., and performed associated mouse viral surgeries in conjunction with S.D.M. and G.E.T.; collected and analyzed behavioral data for mouse IL→MD editing behavioral experiment with GET; Performed TST control experiment in unedited mice; performed mouse histological analysis with K.C. |
| Dalton N. Hughes | Collected and analyzed C57 neurophysiological control data utilized for mouse PYR→PV+ editing experiment. |
| Hannah Schwennesen | Researched gap junction structural/functional properties across animal species, systematically organized findings, selected candidate proteins as substrates for LinCx, and presented recommendations (to RH and KD) with EA, GET, and RB; |
| Stephen D. Mague | Performed all PYR→PV+ and IL→MD (behavioral) mouse viral surgeries in conjunction with K.K.W. and G.E.T.; Performed electrode implantation surgeries in mice with KD. Oversaw mouse PYR→PV+ editing experiment. Supervised mouse IL→MD editing experiments. |
| Daniel Colón-Ramos | Conceived *C elegans* based strategy to assess designer gap junction functional properties; supervised all *C. elegans* experiments; wrote original draft of *in vivo* study results and associated methods with E.W. and A.A.P.; and revised the paper. |
| Rainbo Hultman | Coordinated research of gap junction structural/functional properties across animal species; co-selected fish Cx34.7/Cx35 pair for subsequent protein engineering with KD; Designed Cx34.7/Cx35 mutant library for subsequent cloning; Co-conceptualized framework for screening homotypic then heterotypic interactions in Cx mutants with KD; secured funding and resources; revised the paper. |
| Nenad Bursac | Supervised in vitro studies with KD; secured funding; wrote original draft with E.R., K.C., A.A.P., E.W., D.C.R., and K.D.; and revised the paper. |
| Kafui Dzirasa | Conceived strategy to integrate membrane of two distinct cell types using heterotypically, but not homotypically docking gap junction hemichannels. Co-selected Cx34.7/Cx35 pair for subsequent protein engineering with RCH; Co-conceptualized framework for screening homotypic then heterotypic interactions in Cx mutants with RCH; supervised in vitro studies with NB; supervised computational studies; analyzed data and oversaw all statistical procedures; conceisved all mouse editing experiments; performed electrode implantation surgeries for PYR→PV+ editing experiment and controls with S.D.M. and K.K.W; supervised PYR→PV+ editing experiment; Designed experiment testing $Cx35_{WT}$ with KC, Designed and performed circuit interrogation study using optogenetics with KC; and analyzed data with GET; |

| | |
|---|---|
| | Designed and secured funding and resources; prepared figures; wrote original draft with E.R., K.C., A.A.P., E.W., D.C.R., and N.B.; and revised the paper. |
| **Attribution Process | Each team member outlined their individual contributions across a standard set of domains (conceptualization and methodology, formal analysis, investigation, resources, writing -original draft, writing -review & editing, visualization, Supervision, and Project Administration and Funding Acquisition), and subsequently had the opportunity to edit a summarized attribution description to their satisfaction. Contribution summaries were then shared across all team members. Each team member had the opportunity to raise concerns with regards to any other team member's outlined contributions, and issues that remained unaddressed after additional revisions were subjected to a mediation process led by the lead principal investigator. The assigned authorships and these detailed author contribution descriptions on the initial submitted manuscript deposited in a preprint server reflect the outcome of this process (Ransey et al., 2021). During revisions, additional contributions were made by a subset of the authors. These new attributions were signed off by all authors, and the authorship assignments were revised by the lead principal investigator to reflect each author's composite contributions. Each team member again had the opportunity to raise concerns with regards to any other team member's outlined contributions. |

**Supplementary Table S2, *C. elegans* Strain Table**

| | Genotype | Source | Line # |
|---|---|---|---|
| N2 | *Wild-type* | CGC | |
| DCR3056 | *olaIs17 [Pmod-1::GCaMP6s (25ng/ul) Pttx-3::mCherry (25ng/ul) Punc-122::dsRed (40ng/ul)] I* | Hawk et al. 2018 | |
| DCR6604 | *olaIs23 [Pgcy-8(800)::caPKC-1B (30ng/ul), Pgcy-8(800)::tagRFP (10ng/ul), Punc-122::RFP (30ng/ul)] V; wyIs629 [Pgcy-8(2kb)::GCaMP6s (30ng/ul), Pgcy-8(2kb)::mCherry (5ng/ul), Punc-122:GFP (20ng/ul)] X* | Hawk et al. 2018 | |
| DCR5790 | *olaIs17 [Pmod-1::GCaMP6s (25ng/ul) Pttx-3::mCherry (25ng/ul) Punc-122::dsRed (40ng/ul)] I; olaIs72 [Pelt-7::mCherry (25ng/ul) + Pttx-3::CX36::mCherry (25ng/ul)]* | Hawk et al. 2018 | |
| DCR5793 | *olaIs17 [Pmod-1::GCaMP6s (25ng/ul) Pttx-3::mCherry (25ng/ul) Punc-122::dsRed (40ng/ul)] I; olaIs70 [Pelt-7::GFP (15ng/ul) + Pgcy-8::CX36::mCherry (25ng/ul)]; olaIs72 [Pelt-7::mCherry (25ng/ul) + Pttx-3::CX36::mCherry (25ng/ul)]* | Hawk et al. 2018 | |
| DCR5404 | *olaIs17 [Pmod-1::GCaMP6s (25ng/ul) Pttx-3::mCherry (25ng/ul) Punc-122::dsRed (40ng/ul)] I; olaIs23 [Pgcy-8(800)::caPKC-1B (30ng/ul), Pgcy-8(800)::tagRFP (10ng/ul), Punc-122::RFP (30ng/ul)] V; olaEx3219 [Pgcy-8::CX36::mCherry (25ng/ul); Pttx-3::CX36::mCherry (25ng/ul); Pmyo-3::Red(10ng/ul)]* | Hawk et al. 2018 | |
| DCR8225 | *olaIs17 [Pmod-1::GCaMP6s (25ng/ul) Pttx-3::mCherry (25ng/ul) Punc-122::dsRed (40ng/ul)] I; olaIs23 [Pgcy-8(800)::caPKC-1B (30ng/ul), Pgcy-8(800)::tagRFP (10ng/ul), Punc-122::RFP (30ng/ul)] V 13xOC* | This Paper | |
| DCR8678 | *olaEx5223 [Pgcy-8::CX34.7::GFP; Pttx-3::CX34.7::mCherry; Punc-122::GFP (All 25ng/ul)]* | This Paper | Line 1 |
| DCR8717 | *olaEx5255 [Pgcy-8::CX34.7::GFP; Pttx-3::CX34.7::mCherry; Punc-122::GFP (All 25ng/ul)]* | This Paper | Line 1 |
| DCR8716 | *olaEx5254 [Pgcy-8::CX34.7(E214K, E223K)::GFP; Pttx-3::CX34.7(E214K, E223K)::mCherry; Punc-122::GFP (All 25ng/ul)]* | This Paper | Line 1 |
| DCR8673 | *olaEx5218 [Pgcy-8::CX35(K221E)::GFP; Pttx-3::CX35(K221E)::mCherry; Punc-122::GFP (All 25ng/ul)]* | This Paper | Line 1 |
| DCR8669 | *olaEx5214 [Pgcy-8::CX34.7(E214K, E223K)::GFP; Pttx-3::CX35(K221E)::mCherry; Punc-122::GFP (All 25ng/ul)]* | This Paper | Line 1 |
| DCR8684 | *olaEx5230 [Pgcy-8::CX35::GFP; Pttx-3::CX35::mCherry; Punc-122::GFP (All 25ng/ul)]* | This Paper | Line 2 |
| DCR8719 | *olaEx5256 [Pgcy-8::CX35::GFP; Pttx-3::CX35::mCherry; Punc-122::GFP (All 25ng/ul)]* | This Paper | Line 2 |
| DCR8715 | *olaEx5253 [Pgcy-8::CX34.7(E214K, E223K)::GFP; Pttx-3::CX34.7(E214K, E223K)::mCherry; Punc-122::GFP (All 25ng/ul)]* | This Paper | Line 2 |
| DCR8674 | *olaEx5219 [Pgcy-8::CX35(K221E)::GFP; Pttx-3::CX35(K221E)::mCherry; Punc-122::GFP (All 25ng/ul)]* | This Paper | Line 2 |

| | | | |
|---|---|---|---|
| DCR8676 | *olaEx5221 [Pgcy-8::CX34.7(E214K, E223K)::GFP; Pttx-3::CX35(K221E)::mCherry; Punc-122::GFP (All 25ng/ul)]* | This Paper | Line 2 |
| DCR8685 | *olaEx5231 [Pgcy-8::CX34.7::GFP; Pttx-3::CX34.7::mCherry; Punc-122::GFP (All 25ng/ul)]* | This Paper | Line 3 |
| DCR8720 | *olaEx5257 [Pgcy-8::CX35::GFP; Pttx-3::CX35::mCherry; Punc-122::GFP (All 25ng/ul)]* | This Paper | Line 3 |
| DCR8714 | *olaEx5252 [Pgcy-8::CX34.7(E214K, E223K)::GFP; Pttx-3::CX34.7(E214K, E223K)::mCherry; Punc-122::GFP (All 25ng/ul)]* | This Paper | Line 3 |
| DCR8672 | *olaEx5217 [Pgcy-8::CX35(K221E)::GFP; Pttx-3::CX35(K221E)::mCherry; Punc-122::GFP (All 25ng/ul)]* | This Paper | Line 3 |
| DCR8677 | *olaEx5222 [Pgcy-8::CX34.7(E214K, E223K)::GFP; Pttx-3::CX35(K221E)::mCherry; Punc-122::GFP (All 25ng/ul)]* | This Paper | Line 3 |
| DCR8675 | *olaIs17 [Pmod-1::GCaMP6s (25ng/ul) Pttx-3::mCherry (25ng/ul) Punc-122::dsRed (40ng/ul)] I; olaIs23 [Pgcy-8(800)::caPKC-1B (30ng/ul), Pgcy-8(800)::tagRFP (10ng/ul), Punc-122::RFP (30ng/ul)] V; olaEx5220 [Pgcy-8::CX34.7::GFP; Pttx-3::CX34.7::mCherry; Pelt-7::NLS::mCherry (All 25ng/ul)]* | This Paper | Line 1 |
| DCR8776 | *olaIs17 [Pmod-1::GCaMP6s (25ng/ul) Pttx-3::mCherry (25ng/ul) Punc-122::dsRed (40ng/ul)] I; olaIs23 [Pgcy-8(800)::caPKC-1B (30ng/ul), Pgcy-8(800)::tagRFP (10ng/ul), Punc-122::RFP (30ng/ul)] V; olaEx5287 [Pgcy-8::CX35::GFP; Pttx-3::CX35::mCherry; Punc-122::GFP (All 25ng/ul)]* | This Paper | Line 1 |
| DCR8777 | *olaIs17 [Pmod-1::GCaMP6s (25ng/ul) Pttx-3::mCherry (25ng/ul) Punc-122::dsRed (40ng/ul)] I; olaIs23 [Pgcy-8(800)::caPKC-1B (30ng/ul), Pgcy-8(800)::tagRFP (10ng/ul), Punc-122::RFP (30ng/ul)] V; olaEx5288 [Pgcy-8::CX34.7(E214K, E223K)::GFP; Pttx-3::CX34.7(E214K, E223K)::mCherry; Punc-122::GFP (All 25ng/ul)]* | This Paper | Line 1 |
| DCR8671 | *olaIs17 [Pmod-1::GCaMP6s (25ng/ul) Pttx-3::mCherry (25ng/ul) Punc-122::dsRed (40ng/ul)] I; olaIs23 [Pgcy-8(800)::caPKC-1B (30ng/ul), Pgcy-8(800)::tagRFP (10ng/ul), Punc-122::RFP (30ng/ul)] V; olaEx5216 [Pgcy-8::CX35(K221E)::GFP; Pttx-3::CX35(K221E)::mCherry; Pelt-7::NLS::mCherry (All 25ng/ul)]* | This Paper | Line 1 |
| DCR8670 | *olaIs17 [Pmod-1::GCaMP6s (25ng/ul) Pttx-3::mCherry (25ng/ul) Punc-122::dsRed (40ng/ul)] I; olaIs23 [Pgcy-8(800)::caPKC-1B (30ng/ul), Pgcy-8(800)::tagRFP (10ng/ul), Punc-122::RFP (30ng/ul)] V; olaEx5215 [Pgcy-8::CX34.7(E214K, E223K)::GFP; Pttx-3::CX35(K221E)::mCherry; Pelt-7::NLS::mCherry (All 25ng/ul)]* | This Paper | Line 1 |
| DCR9178 | *olaEx5452 [Pgcy-8::CX34.7(E214K, E223K)::GFP; Pttx-3::CX43::mCherry; Pelt-7::NLS::mCherry (All 25ng/ul)]* | This Paper | Line 1 |
| DCR9179 | *olaEx5453 [Pgcy-8::CX34.7(E214K, E223K)::GFP; Pttx-3::CX43::mCherry; Pelt-7::NLS::mCherry (All 25ng/ul)]* | This Paper | Line 2 |

| | | | |
|---|---|---|---|
| DCR9180 | *olaEx5454 [Pgcy-8::CX34.7(E214K, E223K)::GFP; Pttx-3::CX43::mCherry; Pelt-7::NLS::mCherry (All 25ng/ul)]* | This Paper | Line 3 |
| DCR9181 | *olaIs17 [Pmod-1::GCaMP6s (25ng/ul) Pttx-3::mCherry (25ng/ul) Punc-122::dsRed (40ng/ul)] I; olaIs23 [Pgcy-8(800)::caPKC-1B (30ng/ul), Pgcy-8(800)::tagRFP (10ng/ul), Punc-122::RFP (30ng/ul)] V; olaEx5455 [Pgcy-8::CX34.7(E214K, E223K)::GFP; Pttx-3::CX43::mCherry; Pelt-7::NLS::mCherry (All 25ng/ul)]* | This Paper | Line 1 |
| DCR9182 | *olaEx5456 [Pgcy-8::CX43::GFP; Pttx-3::CX35(K221E)::mCherry; Pelt-7::NLS::mCherry (All 25ng/ul)]* | This Paper | Line 1 |
| DCR9183 | *olaIs17 [Pmod-1::GCaMP6s (25ng/ul) Pttx-3::mCherry (25ng/ul) Punc-122::dsRed (40ng/ul)] I; olaIs23 [Pgcy-8(800)::caPKC-1B (30ng/ul), Pgcy-8(800)::tagRFP (10ng/ul), Punc-122::RFP (30ng/ul)] V; olaEx5457 [Pgcy-8::CX43::GFP; Pttx-3::CX35(K221E)::mCherry; Pelt-7::NLS::mCherry (All 25ng/ul)]* | This Paper | Line 1 |
| DCR9184 | *olaIs17 [Pmod-1::GCaMP6s (25ng/ul) Pttx-3::mCherry (25ng/ul) Punc-122::dsRed (40ng/ul)] I; olaIs72 [Pelt-7::mCherry (25ng/ul) + Pttx-3::CX36::mCherry (25ng/ul)]; olaEx5458 [Pgcy-8::CX34.7(E214K, E223K)::GFP; Pelt-7::NLS::GFP (All 25ng/ul)]* | This Paper | Line 1 |
| DCR9185 | *olaIs17 [Pmod-1::GCaMP6s (25ng/ul) Pttx-3::mCherry (25ng/ul) Punc-122::dsRed (40ng/ul)] I; olaIs72 [Pelt-7::mCherry (25ng/ul) + Pttx-3::CX36::mCherry (25ng/ul)]; olaEx5459 [Pgcy-8::CX34.7(E214K, E223K)::GFP; Pelt-7::NLS::GFP (All 25ng/ul)]* | This Paper | Line 2 |
| DCR9186 | *olaIs17 [Pmod-1::GCaMP6s (25ng/ul) Pttx-3::mCherry (25ng/ul) Punc-122::dsRed (40ng/ul)] I; olaIs72 [Pelt-7::mCherry (25ng/ul) + Pttx-3::CX36::mCherry (25ng/ul)]; olaEx5460 [Pgcy-8::CX34.7(E214K, E223K)::GFP; Pelt-7::NLS::GFP (All 25ng/ul)]* | This Paper | Line 3 |
| DCR7790 | *olaIs17 [Pmod-1::GCaMP6s (25ng/ul) Pttx-3::mCherry (25ng/ul) Punc-122::dsRed (40ng/ul)] I; olaIs70 [Pelt-7::GFP (15ng/ul) + Pgcy-8::CX36::mCherry (25ng/ul)]* | This Paper | |
| DCR9187 | *olaIs17 [Pmod-1::GCaMP6s (25ng/ul) Pttx-3::mCherry (25ng/ul) Punc-122::dsRed (40ng/ul)] I; olaIs70 [Pelt-7::GFP (15ng/ul) + Pgcy-8::CX36::mCherry (25ng/ul)]; olaEx5461 [Pttx-3::CX35(K221E)::mCherry; Pelt-7::NLS::mCherry (All 25ng/ul)]* | This Paper | Line 1 |
| DCR9188 | *olaIs17 [Pmod-1::GCaMP6s (25ng/ul) Pttx-3::mCherry (25ng/ul) Punc-122::dsRed (40ng/ul)] I; olaIs70 [Pelt-7::GFP (15ng/ul) + Pgcy-8::CX36::mCherry (25ng/ul)]; olaEx5462 [Pttx-3::CX35(K221E)::mCherry; Pelt-7::NLS::mCherry (All 25ng/ul)]* | This Paper | Line 2 |

| | | | |
|---|---|---|---|
| DCR9189 | *olaIs17 [Pmod-1::GCaMP6s (25ng/ul) Pttx-3::mCherry (25ng/ul) Punc-122::dsRed (40ng/ul)] I; olaIs70 [Pelt-7::GFP (15ng/ul) + Pgcy-8::CX36::mCherry (25ng/ul)]; olaEx5463 [Pttx-3::CX35(K221E)::mCherry; Pelt-7::NLS::mCherry (All 25ng/ul)]* | This Paper | Line 3 |

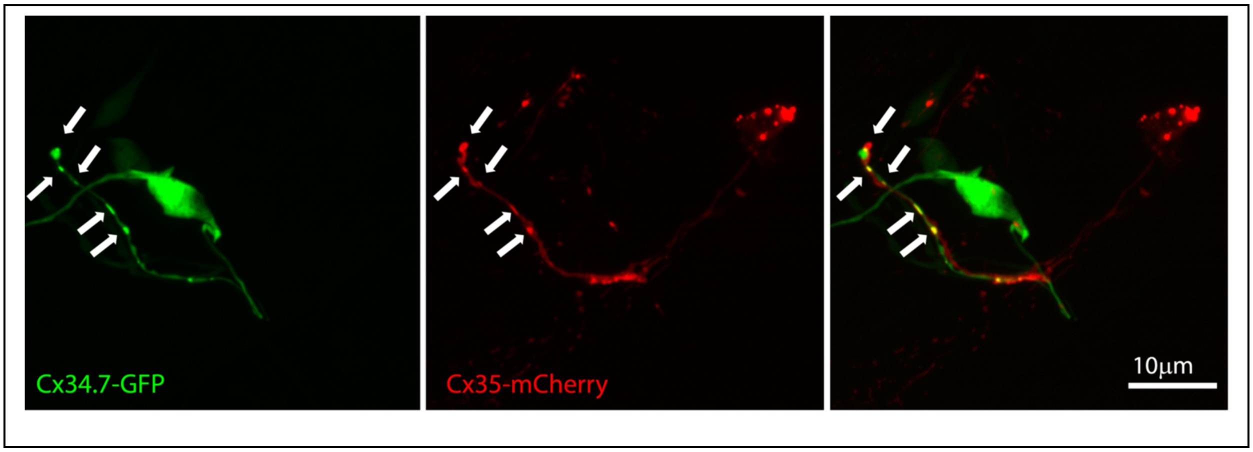


**Supplementary Figure S1: Confocal maximum intensity projections of *C. elegans* expressing the Cx34.7$_{M1}$::Cx35$_{M1}$ pair, related to Fig. 5.** GFP-tagged Cx34.7$_{M1}$ is expressed in the AFD neuron, with puncta along its axon (left). mCherry-tagged Cx35$_{M1}$ is expressed in the AIY neuron, with puncta along its neurite (middle). Composite image showing the colocalizing GFP/mCherry puncta (right). White arrows highlight puncta.

### Cx36 Extracellular Loop 2

| | | | |
|---|---|---|---|
| Homo sapiens | 227– | GLYECNRYPCIKEVECYVSRPTEKTVFLVF | –256 |
| Mus musculus | 227– | GLYECNRYPCIKEVECYVSRPTEKTVFLVF | –256 |
| Macaca mulatta | 227– | GLYECNRYPCIKEVECYVSRPTEKTVFLVF | –256 |
| Callithrix jacchus | 289– | GLYECNRYPCIKEVECYVSRPTEKTVFLVF | –318 |
| Taeniopygia guttata | 196– | AIFECDRYPCVKEVECYVSRPTEKSVFLVF | –225 |
| Danio rerio (Cx34.7) | 209– | GIFECDRYPCLKEVECYVSRPTEKTVFLVF | –238 |
| Danio rerio (Cx35) | 210– | AVYECDRYPCIKDVECYVSRPTEKTVFLVF | –239 |

### Cx43 Extracellular Loop 2

| | | | |
|---|---|---|---|
| Homo sapiens | 171– | LIQWYIYGFSLSAVYTCKRDPCPHQVDCFLSRPTEK | –206 |
| Mus musculus | 171– | LIQWYIYGFSLSAVYTCKRDPCPHQVDCFLSRPTEK | –206 |
| Macaca mulatta | 171– | LIQWYIYGFSLSAVYTCKRDPCPHQVDCFLSRPTEK | –206 |
| Callithrix jacchus | 171– | LIQWYIYGFSLSAVYTCKRDPCPHQVDCFLSRPTEK | –206 |
| Taeniopygia guttata | 171– | LIQWYIYGFSLNAIYTCERDPCPHRVDCFLSRPTEK | –206 |
| Danio rerio | 171– | VIQWYLYGFSLSAVYTCERTPCPHRVDCFLSRPTEK | –206 |

### Cx45 Extracellular Loop 2

| | | | |
|---|---|---|---|
| Homo sapiens | 200– | GFQVHPFYVCSRLPCPHKIDCFI | –222 |
| Mus musculus | 200– | GFQVHPFYVCSRLPCPHKIDCFI | –222 |
| Macaca mulatta | 200– | GFQVHPFYVCSRLPCPHKIDCFI | –222 |
| Callithrix jacchus | 200– | GFQVHPFYVCSRLPCPHKIDCFI | –222 |
| Taeniopygia guttata | 198– | RFEVSPSYVCSRSPCPHTVDCFV | –220 |
| Danio rerio | 196– | GFEVAPSYVCTRSPCPHTVDCFV | –218 |

**Supplementary Figure S2: Sequence alignment of several connexin proteins predicted extracellular loop 2, related to Fig. 6.** Predicted EL2 regions of Cx36 (GJD2), Cx43 (GJA1), and Cx45 (GJC1) for humans and several species broadly utilized in neuroscience research, related to Fig. 6. Identical residues are shown in black, residues that are variable are highlighted with red text. Residues of Cx36 that align to the interaction motif of Cx34.7 and Cx35 are indicated by blue underline. Note zebrafish (Danio rerio) do not have a Cx36 gene, thus the closest homolog, Cx34.7, was used for comparison.

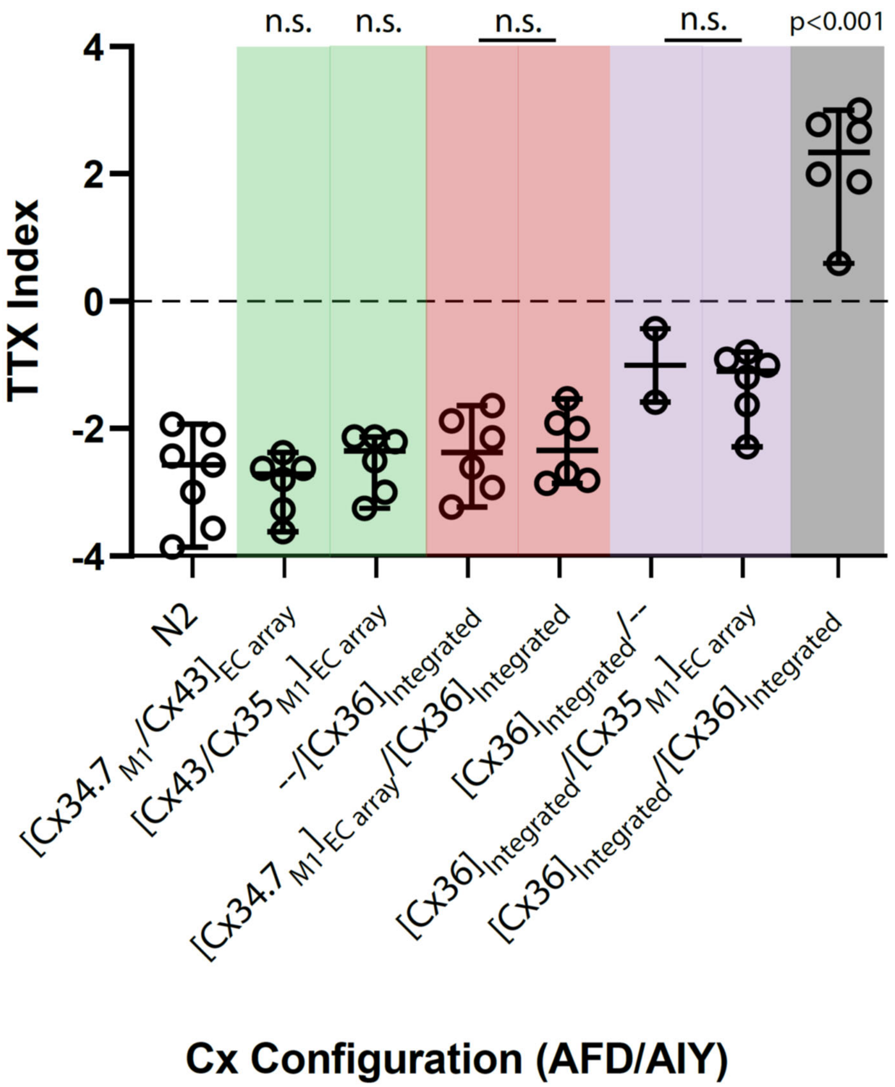


**Supplemental Figure S3: Connexin Cx34.7 and Cx35 do not alter *C. elegans* behavior when expressed counterparts to connexins endogenous to the human brain.** We tested Cx34.7$_{M1}$ and Cx35$_{M1}$ hemichannels against Cx36 and Cx43. Worms with integrated arrays were used to test the mutant channels against Cx36. TTX Index (Y axis) refers to the thermotaxis index, calculated as previously described (Hawk et al., 2018). In the X-axis, N2 refers to the wild type *C. elegans* animals expressing no arrays. The other groups describe the expression of specific arrays in an AFD (presynaptic)/AIY (postsynaptic) configuration. "EC array" refers to extrachromosomal arrays, while "Integrated" refers to integrated arrays. Experimental worms were directly compared against integrated Cx36 controls, where Cx34.7$_{M1}$ or Cx35$_{M1}$ expression patterns mirrored our original experimental configuration. Worms expressing extrachromosomal arrays in AFD or AIY were used to test Cx43. C. elegans expressing Cx34.7$_{M1}$/Cx36, Cx34.7$_{M1}$/Cx43, Cx36/Cx35$_{M1}$ or Cx43/Cx35$_{M1}$ in AFD/AIY all continued to migrate towards cold temperatures ($F_{7,10.67}$= 19.29; P<0.0001 using Welch one-way ANOVA followed by Dunnett's T3 multiple comparisons; p<0.001), and no differences were observed across experimental pairs (P>0.05; highlighted by red or purple), indicating they were not forming functional electrical synapses.

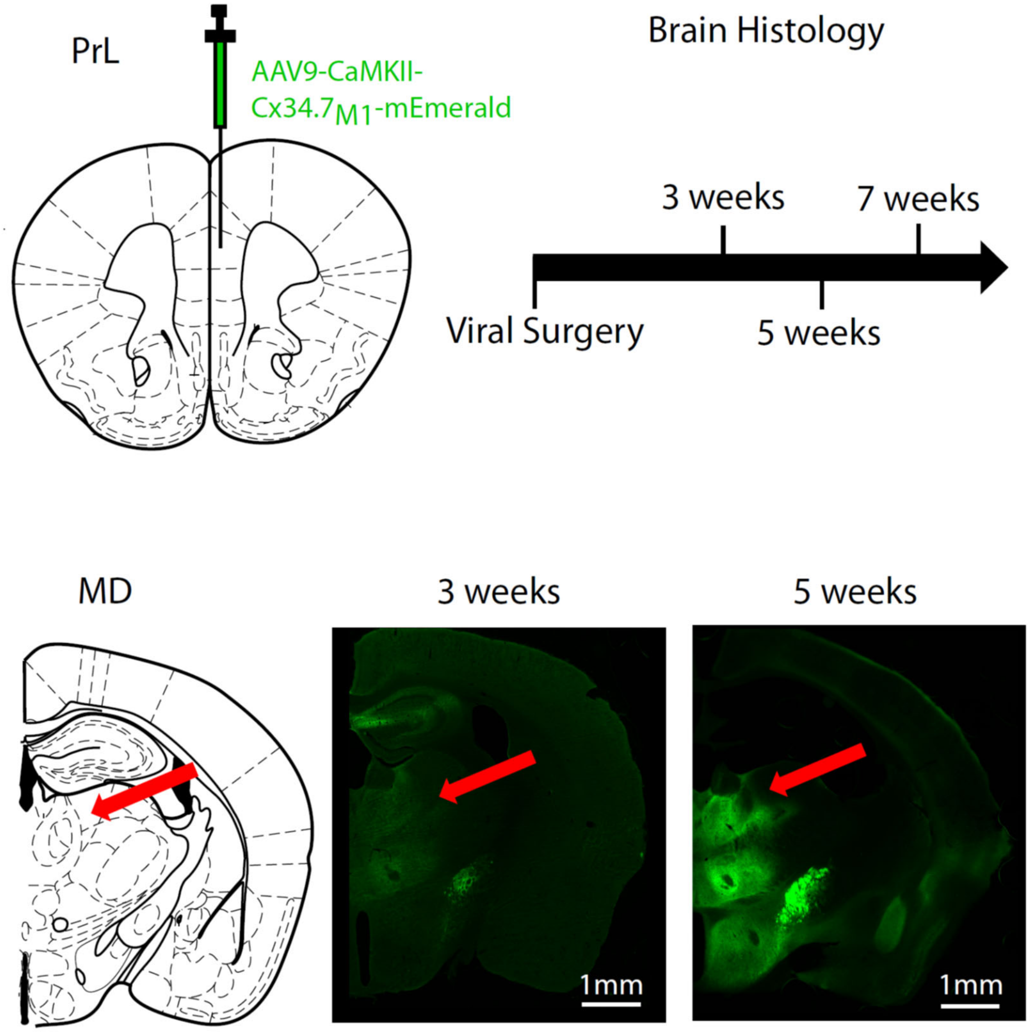


| | 3 weeks | 5 weeks | 7 weeks |
|---|---|---|---|
| Dorsal Striatum | X | X | X |
| Nucleus Accumens | | X | X |
| Basolateral Amygdala | | X | X |
| Medial Dosal Thalamus | | X | X |
| Ventral Tegmental Area | | | X |

**Supplemental Figure S4: Time course of pronounced Cx34.7 expression and trafficking in prelimbic cortex soma and terminals, related for Fig 6.** Five C57BL/6J mice were infected with AAV9-CaMKII-Cx34.$7_{M1}$-mEmerald in Prelimbic cortex. Histology was performed 3 weeks (N=1 mouse), 5 weeks (N=2 mice), or 7 weeks (N=2 mice) after viral surgeries.

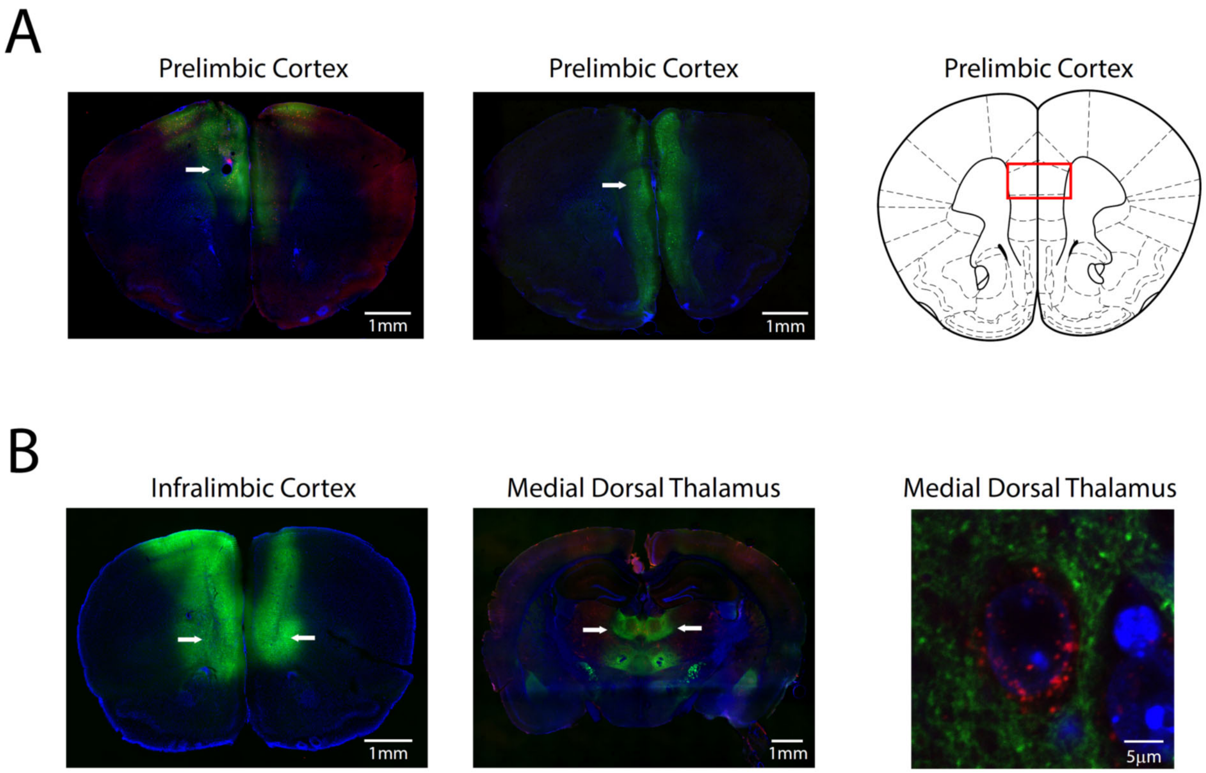


**Supplemental Figure S5: Representative histological images for mouse editing experiments, related to Fig. 5 and 6. A)** Representative histological image of prelimbic cortex in A) Cx34.7$_{M1}$/ Cx35$_{M1}$ infected mouse (left) and a Cx34.7$_{M1}$/ Cx34.7$_{M1}$ control (middle). White arrows highlight electrode tracks. For mice included in the prelimbic cortex editing experiment, individual electrode wires were distributed within the area highlighted by the red rectangle. **B)** Histological image of mouse injected with AAV9-CaMKII-Cx34.7$_{M1}$-mEmerald in infralimbic cortex (left). White arrows highlight viral injection tracks. Image showing medial dorsal thalamus in mouse injected with AAV9-CaMKII-Cx34.7$_{M1}$-mEmerald in infralimbic cortex and AAV9-CaMKII-Cx35$_{M1}$-mApple in medial dorsal thalamus (middle). Composite confocal image of medial dorsal thalamus showing expression of Cx35$_{M1}$-mApple (red) at the soma of a cell stained with DAPI (blue) and Cx34.7$_{M1}$-mEmerald (green) at IL nerve terminals.

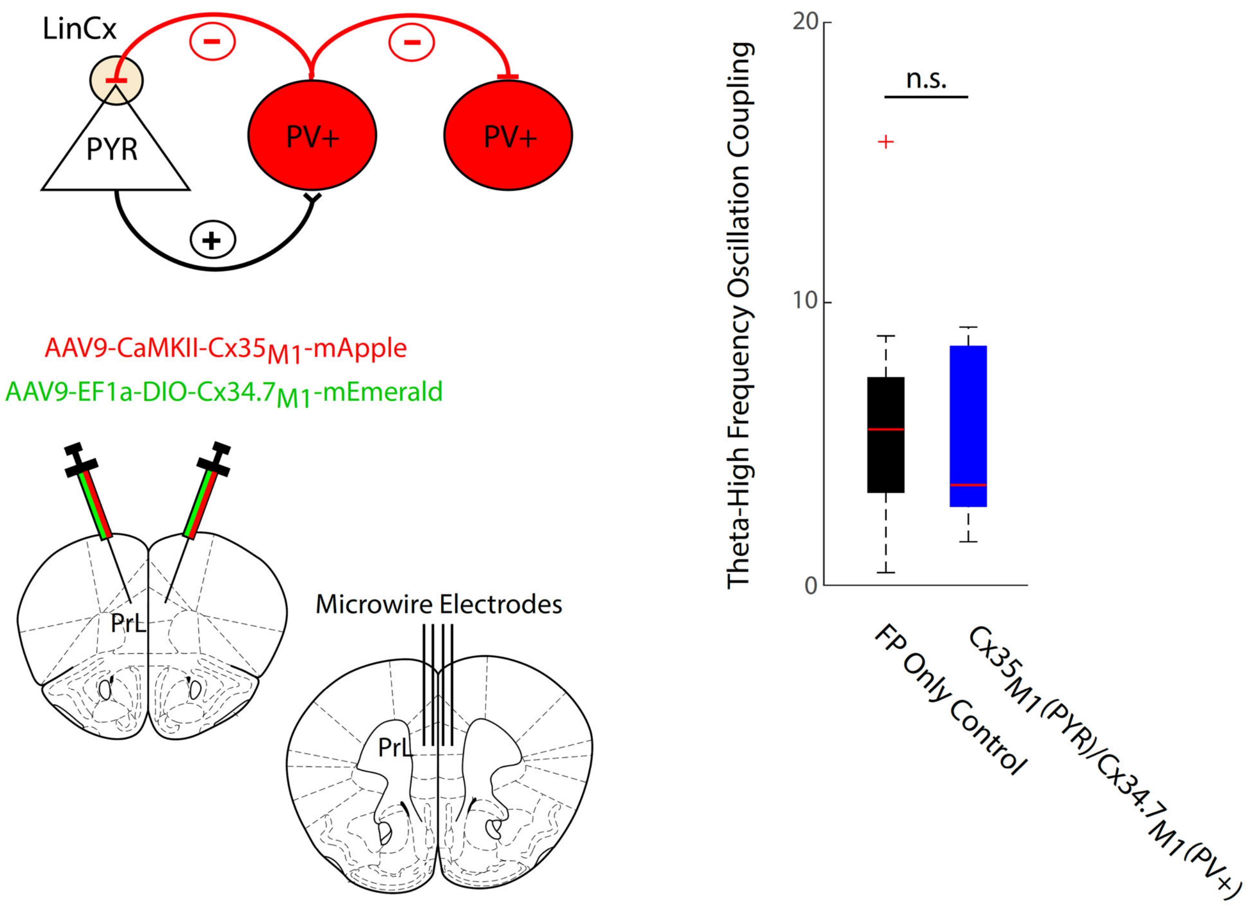


**Supplemental Figure S6: Editing parvalbumin expressing interneuron→pyramidal neuron circuit with LinCx does not enhance theta-high frequency oscillation coupling.** The experimental schematic for viral infection and electrode implantation is shown to the left. No differences were observed between controls and the mice expressing $Cx35_{M1}$/$Cx34.7_{M1}$ to target pyramidal (PYR) and parvalbumin-expressing interneurons (PV+; U= 125, P=0.92 using two tailed rank-sum test).

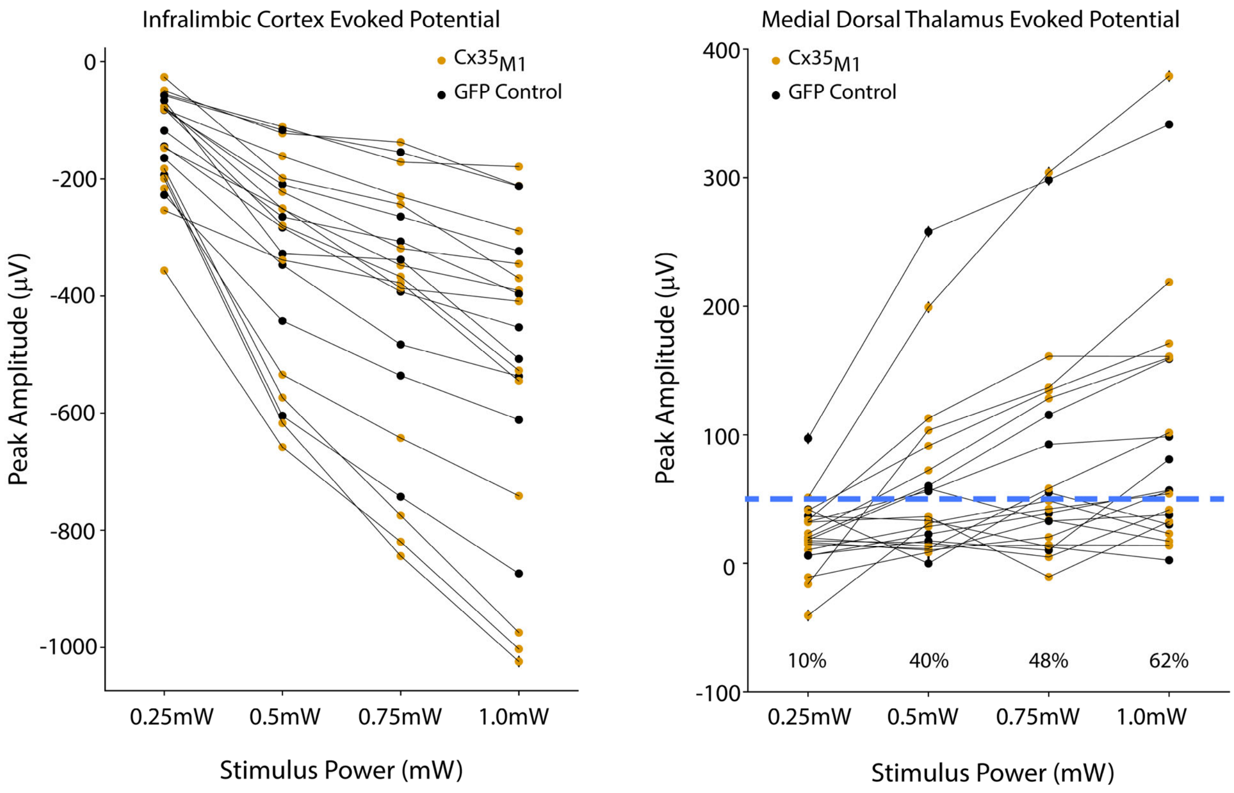


**Supplemental Figure S7: Cortical and thalamic potentials evoked by IL stimulation during first experimental session.** Peak cortical amplitudes within 10ms of stimulation are shown to the left. Responses were averaged across microwires. Peak thalamic amplitudes within 25ms are shown the right. The percentage of mice that showed thalamic evoked responses above 50mV is shown for each light intensity. N=21 total mice.

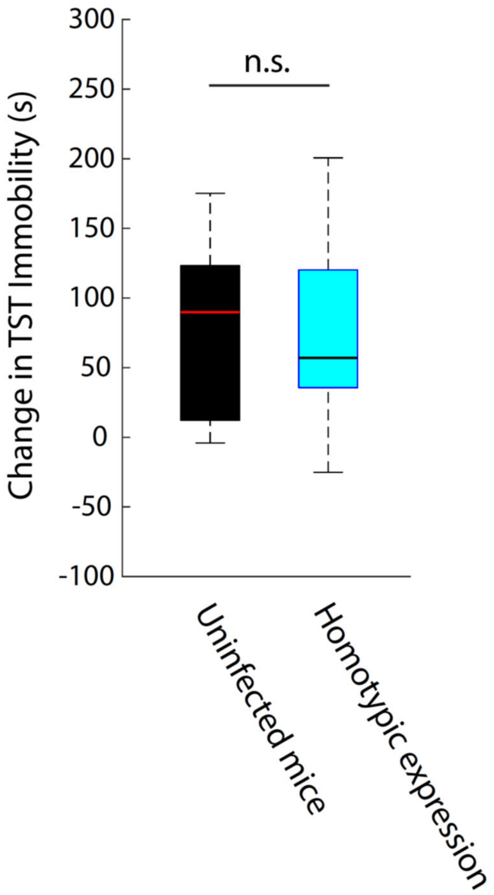


**Supplemental Figure S8: Homotypic expression of Cx34.7$_{M1}$ or Cx35$_{M1}$ across the IL→MD circuit does not impact stress inducted behavioral adaptation ($T_{29}$=0.16, P=0.87 using two tailed t-test).**

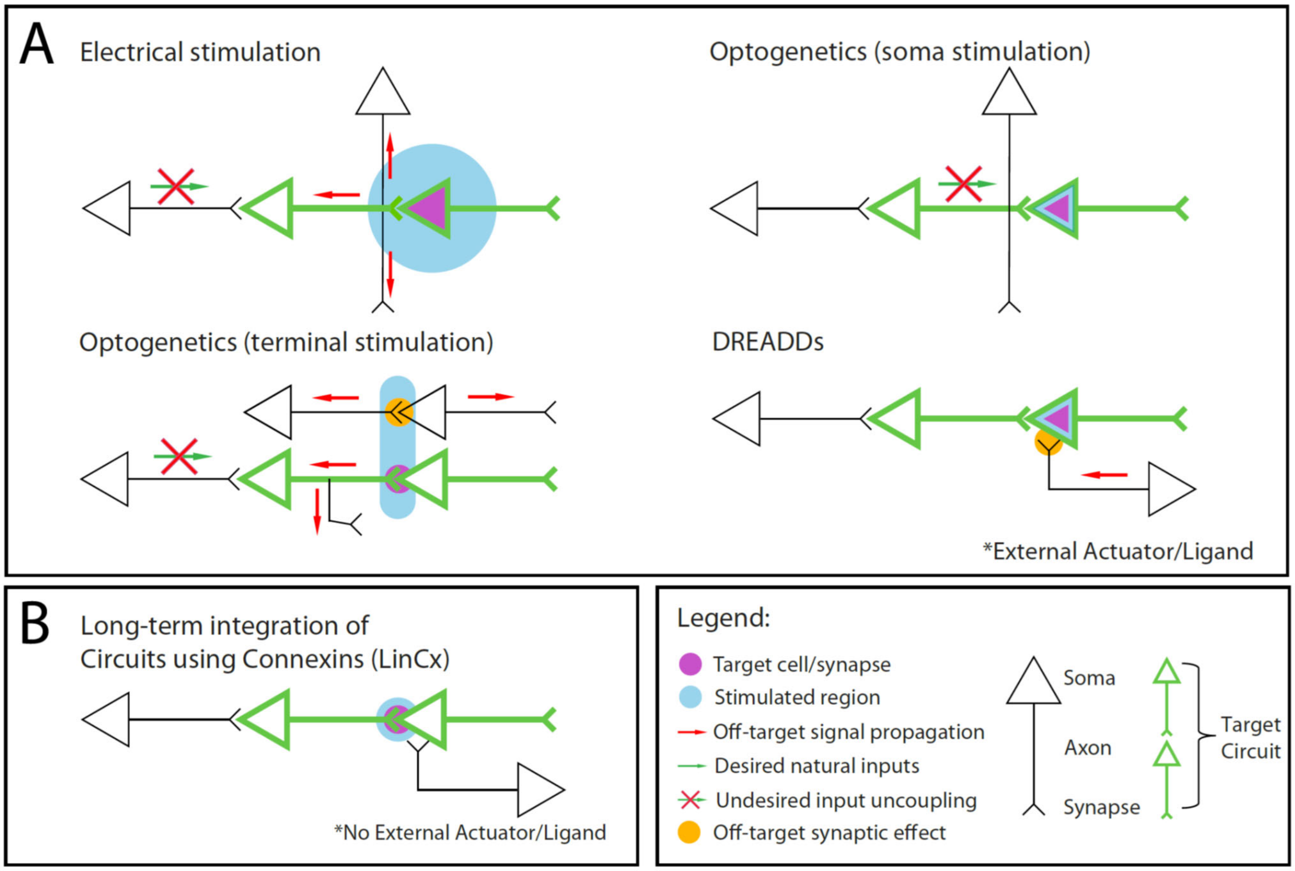


**Supplemental Figure S9. Complementary utility of LinCx compared to widely adopted neural modulation approaches in selectively targeting a spatially and cell-type pair defined circuit. A)** Electrical stimulation activates many cell types and pass-through fibers within a tissue volume. Stimulation also modulates cellular activity independent of the context of input fibers, and potentially drives retrograde activation of cellular inputs. Open-loop optogenetic stimulation of neuronal soma modulates cell-type specific activity independent of the context of input fibers. Optogenetic terminal stimulation enables modulation of connections between brain regions, but potentially activates multiple circuits defined by distinct post-synaptic cell types, and potentially drives retrograde activation of target axons. DREADDs modulate target cells, but potentially modulate the response of target cells to their input fibers. **B)** LinCx enables selective targeting based on a cell-type defined pair and selective modulation of the post-synaptic cell-type based on the activity context of the pre-synaptic cell.

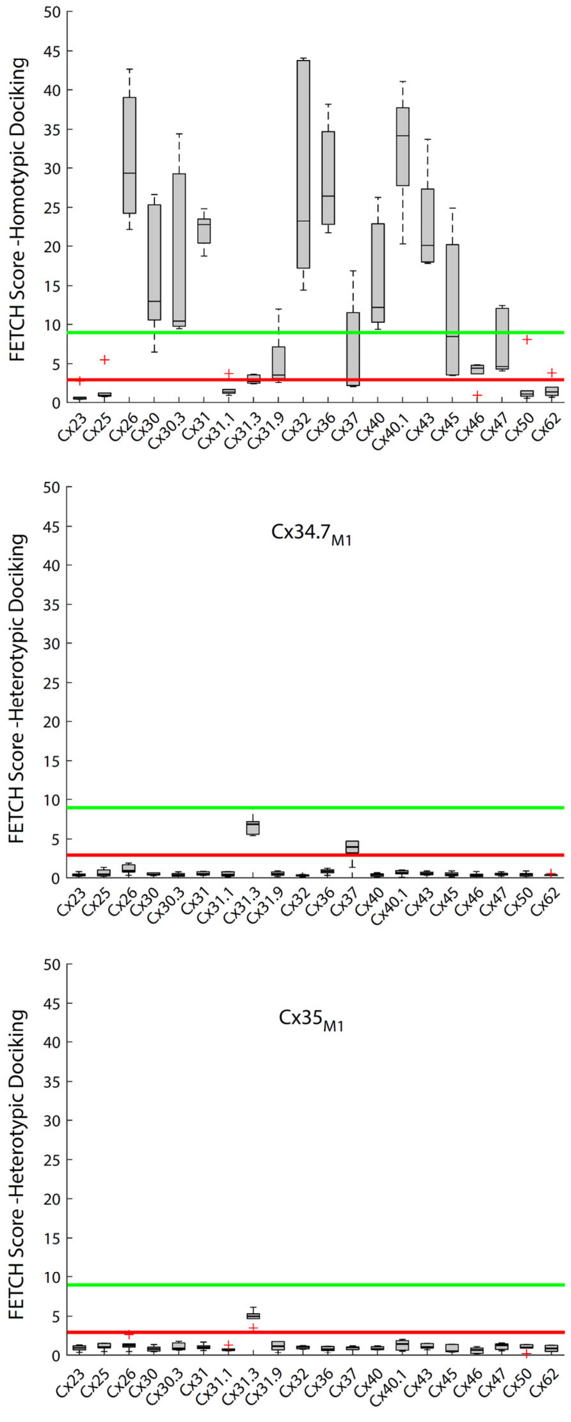


**Supplemental Figure S10. Docking compatibility of Cx34.7$_{M1}$ and Cx35$_{M1}$ with human connexins.** We used FETCH to quantify the homotypic docking characteristics of 20 human connexin isoforms (top). We then characterized the heterotypic docking characteristic of our mutant proteins with the 20 endogenous isoforms (middle and bottom). The red and green lines correspond to the upper and lower limits of non-docking and docking population shown in Fig 1F, respectively. Notably, expression of our Cx59 constructs appeared to promote cell death and was thus excluded from our studies.
